# Massively multiplex single-molecule oligonucleosome footprinting

**DOI:** 10.1101/2020.05.20.105379

**Authors:** Nour J Abdulhay, Colin P McNally, Laura J Hsieh, Sivakanthan Kasinathan, Aidan Keith, Laurel S Estes, Mehran Karimzadeh, Jason G Underwood, Hani Goodarzi, Geeta J Narlikar, Vijay Ramani

## Abstract

Our understanding of the beads-on-a-string arrangement of nucleosomes has been built largely on high-resolution sequence-agnostic imaging methods and sequence-resolved bulk biochemical techniques. To bridge the divide between these approaches, we present the single-molecule adenine methylated oligonucleosome sequencing assay (SAMOSA). SAMOSA is a high-throughput single-molecule sequencing method that combines adenine methyltransferase footprinting and single-molecule real-time DNA sequencing to natively and nondestructively measure nucleosome positions on individual chromatin fibres. SAMOSA data allows unbiased classification of single-molecular ‘states’ of nucleosome occupancy on individual chromatin fibres. We leverage this to estimate nucleosome regularity and spacing on single chromatin fibres genome-wide, at predicted transcription factor binding motifs, and across both active and silent human epigenomic domains. Our analyses suggest that chromatin is comprised of a diverse array of both regular and irregular single-molecular oligonucleosome patterns that differ subtly in their relative abundance across epigenomic domains. This irregularity is particularly striking in constitutive heterochromatin, which has typically been viewed as a conformationally static entity. Our proof-of-concept study provides a powerful new methodology for studying nucleosome organization at a previously intractable resolution, and offers up new avenues for modeling and visualizing higher-order chromatin structure.

**1-sentence summary:** High-throughput single-molecule real-time footprinting of chromatin arrays reveals heterogeneous patterns of oligonucleosome occupancy.

## INTRODUCTION

The nucleosome is the atomic unit of chromatin. Nucleosomes passively and actively template the majority of nuclear interactions essential to life by determining target site access for transcription factors^1^, bookmarking active and repressed chromosomal compartments via post-translational modifications^2^, and safeguarding the genome from mutational agents^3^. Our earliest views of the beads-on-a-string arrangement of chromatin derived from classical electron micrographs of chromatin fibres^4^, which have since been followed by both light^5^ and electron microscopy^6,7^ studies of *in vitro*-assembled and *in vivo* chromatin. In parallel, complementary biochemical methods using nucleolytic cleavage have successfully mapped the subunit architecture of chromatin structure at high resolution. These cleavage-based approaches can be stratified into those that focus primarily on chromatin accessibility^8^ (*i.e*. measuring ‘competent’ active chromatin^9^), and those that survey nucleosomal structure uniformly across active and inactive genomic compartments. Understanding links between chromatin and gene regulation requires sensitive methods in all three of these broad categories: in this study we advance our capabilities in the third, focusing on a novel method to map oligonucleosomal structures genome-wide.

Nucleolytic methods for studying nucleosome positioning have historically used cleavage reagents (*e.g*. dimethyl sulphate^10^, hydroxyl radicals^11^, nucleases^12^) followed by gel electrophoresis and / or Southern blotting to map the abundance, accessibility, and nucleosome repeat lengths (NRLs) of chromatin fibers^13^. More recently, these methods have been coupled to high-throughput short-read sequencing^14^, enabling genome-wide measurement of average nucleosome positions. While powerful, all of these methods share key limitations: measurement of individual protein-DNA interactions inherently requires destruction of the chromatin fibre and averaging of signal across many short molecules. These limitations extend even to single-molecule methyltransferase-based approaches^15–17^, which have their own biases (*e.g*. CpG / GpC bias; presence of endogenous m^5^dC in mammals; DNA damage due to bisulphite conversion), and are still subject to the short-length biases of Illumina sequencers. While single-cell^18,19^ and long-read single-molecule^20^ genomic approaches have captured some of this lost contextual information, single-cell data are generally sparse and single-molecule Array-seq data must be averaged over multiple molecules. Ultimately, these limitations have hindered our understanding of how combinations of ‘oligonucleosomal patterns’ (*i.e*. discrete states of nucleosome positioning and regularity on single DNA molecules) give rise to active and silent chromosomal domains.

The advent of third-generation (*i.e*. high-throughput, long-read) sequencing offers a potential solution to many of these issues^21^. Here, we demonstrate <u>S</u>ingle-molecule <u>A</u>denine <u>M</u>ethylated <u>O</u>ligonucleosome <u>S</u>equencing <u>A</u>ssay (SAMOSA), a method that combines adenine methyltransferase footprinting of nucleosomes with base modification detection on the PacBio single-molecule real time sequencer^22^ to measure nucleosome positions on single chromatin templates. We first present proof-of-concept of SAMOSA using gold-standard *in vitro* assembled chromatin fibres, demonstrating that our approach captures single-molecule nucleosome positioning at high-resolution. We next apply SAMOSA to oligonucleosomes derived from K562 cells to profile single-molecule nucleosome positioning genome-wide. Our data enables unbiased classification of oligonucleosomal patterns across both euchromatic and heterochromatic domains. These patterns are influenced by multiple epigenomic phenomena, including the presence of predicted transcription factor binding motifs and post-translational histone modifications. Consistent with estimates from previous studies, our approach reveals enrichment for long, regular chromatin arrays in actively elongating chromatin, and highly accessible, disordered arrays at active promoters and enhancers. Surprisingly, we also observe a large amount of heterogeneity within constitutive heterochromatin domains, with both mappable H3K9me3-decorated regions and human major satellite sequences harboring a mixture of irregular and short-repeat-length oliognucleosome types. Our study provides a proof-of-concept framework for studying chromatin at single-molecule resolution while suggesting a highly dynamic nucleosome-DNA interface across chromatin sub-compartments.

## RESULTS

### Single-molecule real-time sequencing of adenine-methylated chromatin captures nucleosome footprints

Existing methyltransferase accessibility assays either rely on bisulfite conversion^15–17^ or use the Oxford Nanopore platform to detect DNA modifications^23–25^. We hypothesized that high-accuracy PacBio single-molecule real-time sequencing could detect m^6^dA deposited on chromatin templates to natively measure nucleosome positioning. To test this hypothesis, we used the nonspecific adenine methyltransferase EcoGII^26^ to footprint nonanucleosomal chromatin arrays generated through salt-gradient dialysis (**Supplementary Figure 1**), using template DNA containing 9 tandem repetitive copies of the Widom 601 nucleosome positioning sequence^27^ separated by ~46 basepairs (bp) of linker sequence followed by ~450 bp of sequence without any known intrinsic affinity for nucleosomes. After purifying DNA, polishing resulting ends, and ligating on barcoded SMRTBell adaptors, we subjected libraries to sequencing on PacBio Sequel or Sequel II flow cells, using unmethylated DNA and methylated naked DNA as controls (**Figure 1A**). After filtering low quality reads, we analyzed a total of 33,594 single molecules across all three conditions. Across both platforms, we observed higher average interpulse duration (IPD) in samples exposed to methyltransferase, consistent with a rolling circle polymerase ‘pausing’ at methylated adenine residues in template DNA (**Supplementary Figure 2**). Further inspection of footprinted chromatin samples sequenced on either platform revealed strong specificity for altered IPD values only at thymines falling outside Widom 601 repeat sequences, in contrast with fully methylated naked template and unmethylated controls (**Supplementary Figure 3A,B**). These results suggest that the PacBio platform can natively detect ectopic m^6^dA added to chromatinized templates.

**Figure 1:**
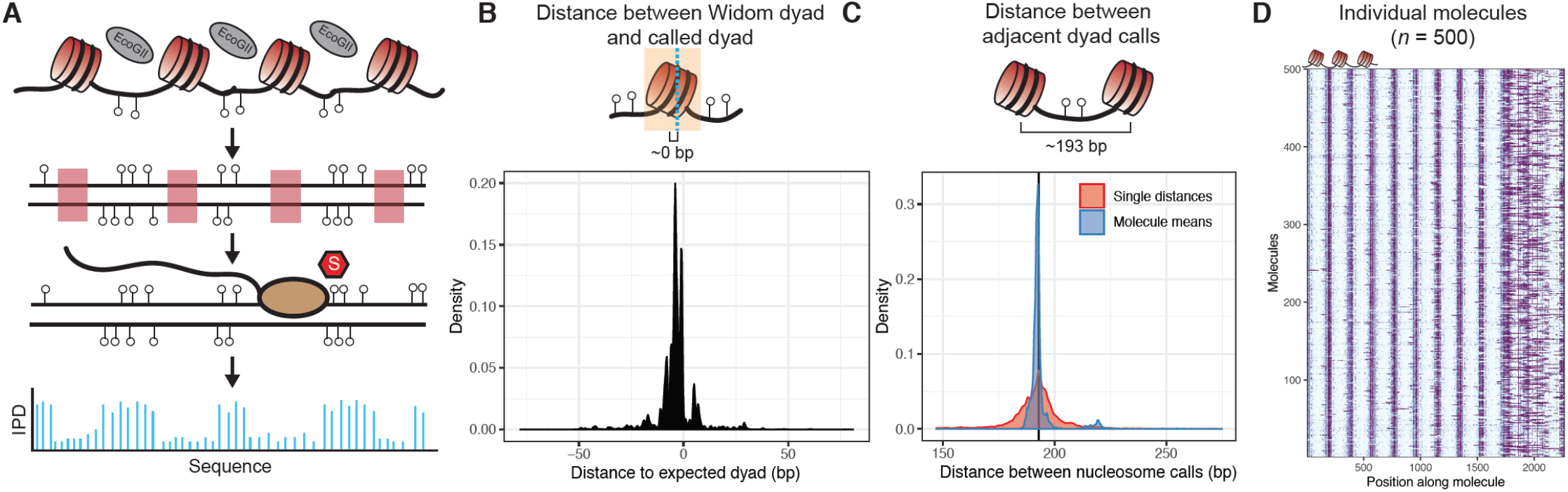
Overview of the single-molecule adenine methylated oligonucleosome sequencing assay (SAMOSA). **A.)** In the SAMOSA assay, chromatin is methylated using the nonspecific EcoGII methyltransferase, DNA is purified, and then subjected to sequencing on the PacBio platform. Modified adenine residues are natively detected during SMRT sequencing due to polymerase pausing, leading to an altered interpulse duration at modified residues. **B.)** SAMOSA data can be used to accurately infer nucleosome dyad positions given a strong positioning sequence. Shown are the distributions of called dyad positions with respect to the known Widom 601 dyad. Called dyads fall within a few nucleotides of the expected dyad position (median ± median absolute deviation [MAD] = 4 ± 2.97 bp). **C.)** SAMOSA data accurately recapitulates the known nucleosome repeat lengths (NRL) of *in vitro* assembled chromatin fibres. Called NRLs are strongly concordant with the expected 193 repeat length (pairwise distance between adjacent dyads median ± MAD = 193 ± 7.40 bp; single-molecule averaged repeat length median ± MAD = 192 ± 1.30 bp). **D.)** Expected nucleosome footprints in SAMOSA data can be visually detected with single-molecule resolution (*n* = 500 sampled footprinted chromatin molecules).

We next developed a computational approach to assign a posterior probability describing the likelihood that an A/T basepair is methylated given IPD signals found within the same molecule (*i.e*. ‘modification probability’). We then paired this approach with a simple peak-calling strategy to approximate nucleosomal dyad positions. To benchmark this pipeline, we first calculated the distance between called nucleosome dyads and expected 601 dyad positions (**Figure 1B**). Observed dyads were highly concordant with expected positions (median ± median absolute deviation [MAD] = 4 ± 2.97 bp), consistent with our data accurately capturing the expected 601 dyad. We next calculated the expected distances between nucleosomes given our dyad callset (*i.e*. a computationally-defined nucleosome repeat length [NRL]; **Figure 1C**). Compared with the expected repeat length of 193 bp, our calculated results were similarly accurate at both two-dyad resolution (pairwise distance between adjacent dyads; median ± MAD = 193 ± 7.40 bp) and averaged single-molecule resolution (median ± MAD = 192 ± 1.30 bp). Both of these measurements were qualitatively uniform across all molecules, independent of the positions of individual nucleosomes along individual array molecules (**Supplementary Figure 4**). Finally, we directly visualized the modification probabilities of individual sequenced chromatin molecules and observed that modification patterns occurred in expected linker sequences (**Figure 1D**), and not in unmethylated or fully methylated control samples (**Supplementary Figure 5A,B**). These results demonstrate that EcoGII footprinting is specific for unprotected DNA and that kinetic deviations observed in the data are not simply the result of primary sequence biases in the template itself. We hereafter refer to this approach as the <u>S</u>ingle-molecule <u>A</u>denine <u>M</u>ethylated <u>O</u>ligonucleosome <u>S</u>equencing <u>A</u>ssay (SAMOSA).

### SAMOSA captures regular nucleosome-DNA interactions *in vivo* through nuclease-cleavage and adenine-methylation simultaneously

Having shown that SAMOSA can footprint *in vitro* assembled chromatin fibres, we sought to apply our approach to oligonucleosomal fragments from living cells. Multiple prior studies have suggested that a light micrococcal nuclease (MNase) digest followed by disruption of the nuclear envelope and overnight dialysis can be used to gently liberate oligonucleosomes into solution without dramatically perturbing nucleosomal structure^28–30^. After lightly digesting and solubilizing oligonucleosomes from human K562 nuclei, we methylated chromatin with EcoGII and sequenced methylated molecules on the Sequel II platform (*n* = 1,855,316 molecules total; **Figure 2A**). As controls, we also shallowly sequenced deproteinated K562 oligonucleosomal DNA, and deproteinated oligonucleosomal DNA methylated with the EcoGII enzyme.

**Figure 2:**
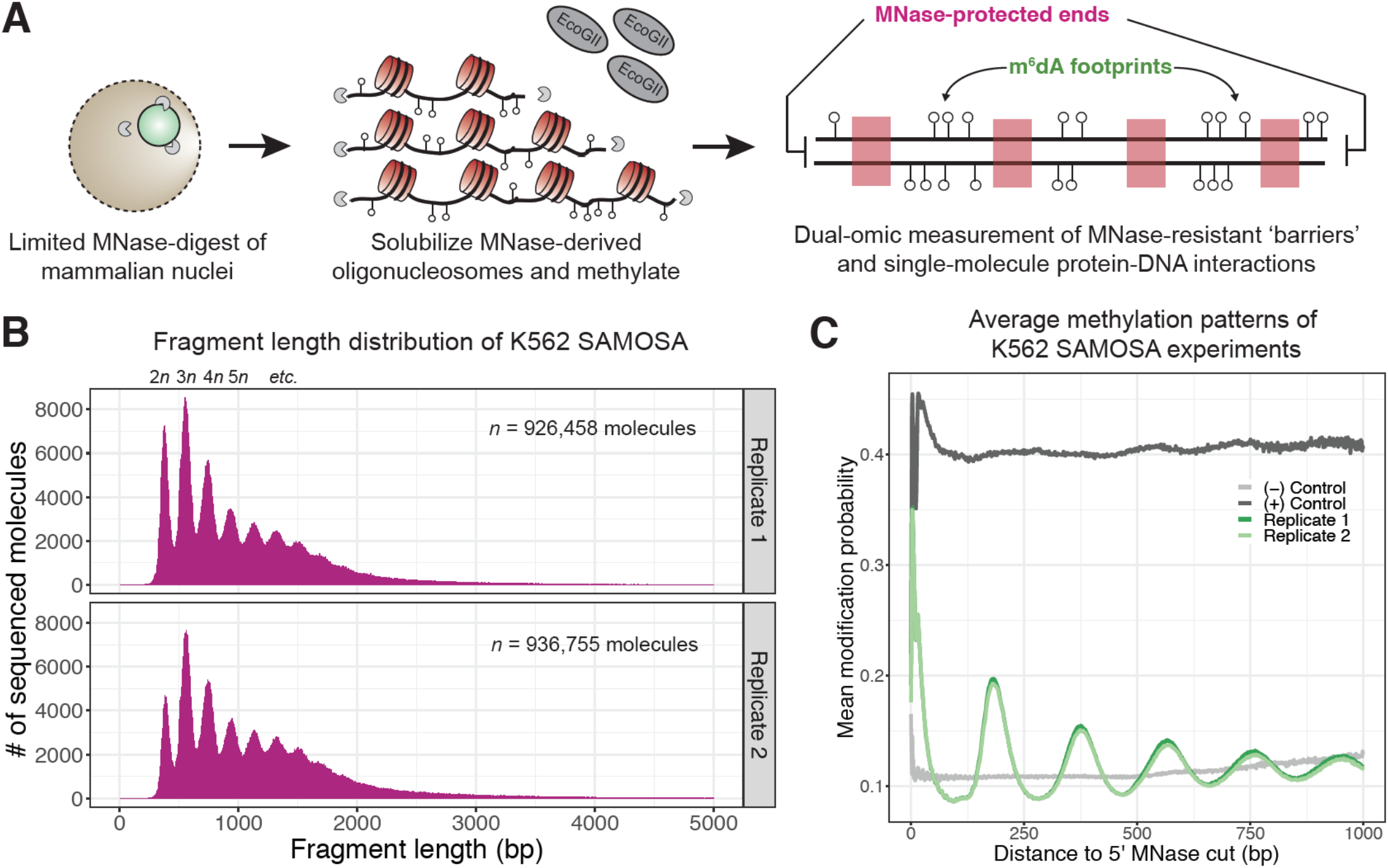
*In vivo* SAMOSA captures oligonucleosome structure by combining MNase digestion of chromatin with adenine methylation footprinting. **A.)** An overview of the *in vivo* SAMOSA protocol: oligonucleosomes are gently solubilized from nuclei using micrococcal nuclease and fusogenic lipid treatment. Resulting oligonucleosomes are footprinted using the EcoGII enzyme and sequencing on the PacBio platform. Each sequencing molecules captures two orthogonal biological signals: MNase cuts that capture ‘barrier’ protein-DNA interactions, and m^6^dA methylation protein-DNA footprints. **B.)** Fragment length distributions for *in vivo* SAMOSA data reveal expected oligonucleosomal laddering (bin size = 5 bp). **C.)** Averaged modification probabilities from SAMOSA experiments demonstrate the ability to mark nucleosome-DNA interactions directly via methylation. Modification patterns seen in the chromatin sample are *not* seen in unmethylated oligonucleosomal DNA or fully methylated K562 oligonucleosomal DNA.

*In vivo* SAMOSA has several advantages compared to existing MNase- or methyltransferase-based genomic approaches. Our approach combines MNase-derived cuts flanking each fragment with methyltransferase footprinting of nucleosomes. MNase cuts mark the boundary of genomic ‘barrier’ elements like nucleosomes; accordingly, fragment length distributions from *in vivo* SAMOSA data display patterns emblematic of bulk nucleosomal array regularity (**Figure 2B**). Modification patterns of sequenced molecules then capture nucleosome-positioning information at single-molecule resolution; this is evident in single-molecule averages of modification probability in chromatin samples with respect to fully methylated and unmethylated controls (**Figure 2C**). While previous approaches for studying nucleosome regularity may capture each of the former information types, this method is, to our knowledge, the first “multi-omic” single-molecule assay that simultaneously captures the positioning of protein-DNA interactions through nucleolytic cleavage, and (through DNA methylation) the positioning of proximal protein-DNA interactions on the same single-molecule.

Our approach also differs from recent long-read^23,24,31^ surveys of the epigenome in that our experiments capture both active and repressed epigenomic domains. Comparing coverage enrichment of specific ChromHMM^32^ labels in our dataset to shotgun PacBio sequencing of a human genome, ENCODE K562 DNase I coverage^33^, and published K562 MNase-seq coverage^34^, we observe notably increased representation of heterochromatic regions of the genome with respect to DNase I data (**Supplementary Figure 6)**, albeit with different biases not seen in shotgun sequencing (likely owing to a combination of our solubilization protocol and MNase fragmentation). Finally, our approach relies on PacBio sequencing, which at the moment has demonstrably higher accuracy^35^ than Nanopore based approaches, and does not use the bisulfite conversion required by Illumina-based methods to detect modifications.

### SAMOSA enables unbiased classification of chromatin fibres on the basis of regularity and nucleosome repeat length

The relative abundance and diversity of oligonucleosome patterns across the human genome remains unknown. Given the single-molecule nature of SAMOSA, we speculated that our data could be paired with a state-of-the-art community detection algorithm to systematically cluster footprinted molecules on the basis of single-molecule nucleosome regularity and NRL (*i.e*. ‘oligonucleosome patterns’). To ease detection of signal regularity on single molecules, we computed autocorrelograms for each molecule in our dataset ≥ 500 bp in length, and subjected resulting values to unsupervised Leiden clustering^36^. Cluster sizes varied considerably, but were consistent across both replicates, with each cluster containing 6.54% (Cluster 4) - 20.1% (Cluster 1) of all molecules (**Figure 3A**). The resulting seven clusters (**Supplementary Figure 7A**) capture the spectrum of oligonucleosome patterning genome-wide, stratifying the genome by both NRL and array regularity. Accounting for the coverage biases presented above, the measurements shown in **Figure 3A** provide a rough estimate of the equilibrium composition of the genome with respect to these patterns.

**Figure 3:**
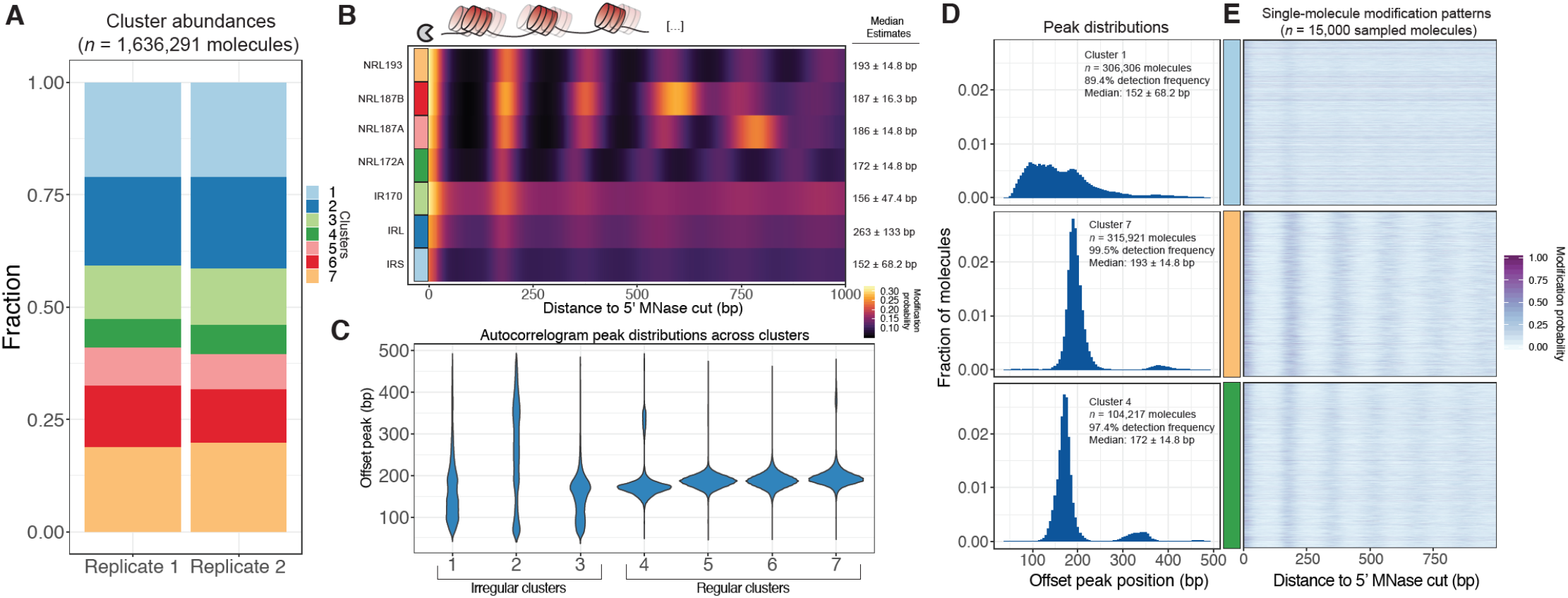
SAMOSA reveals distribution of oligonucleosome patterns genome-wide. **A.)** Stacked bar chart representation of the contribution of each cluster to overall signal across two replicate experiments in K562 cells. **B.)** Average modification probability as a function of sequene for each of the seven defined clusters. *Left:* Manually annoted cluster names based on NRL estimates computed by calling peaks on single-molecule autocorrelograms; *Right:* Median and median absolute deviation for single-molecule NRL estimates determined for each cluster. **C.)** Violin plot representation of the distributions of single-molecule NRL estimates for each cluster. Clusters can be separated into three ‘irregular’ and four ‘regular’ groups of oligonucleosomes. **D.)** Histogram of single-molecule NRL estimates for Clusters 1, 4, and 7, along with **E.)** 5,000 randomly sampled molecules from each cluster.

The diversity in nucleosome regularity and repeat length across these clusters is visually apparent when inspecting average modification probabilities of the 5’ 1000 bp of each cluster (**Figure 3B**). To better annotate each of these clusters, we characterized each with respect to methylation extent and distributions of computed single-molecule NRLs. We first inspected the average modification probabilities of each molecule across clusters, finding that these averages were largely invariant (**Supplementary Figure 7B**). This suggests that our clustering approach does not simply classify oligonucleosomes based on the amount of methylation on each molecule. We next estimated within-cluster heterogeneity in single-molecule NRLs using a simple peak-calling approach. We scanned each autocorrelogram for secondary peaks, and annotated the location of each peak to compute an estimated NRL. We then visualized these distributions as violin plots for each cluster (**Figure 3C**). Our data broadly fall into two categories: irregular clusters made up of molecules spanning multiple NRLs and lacking a strong regular periodicity, and highly regular clusters with defined single-molecule NRLs ranging from ~172 bp (*i.e*. chromatosome plus 5 bp DNA) to >200 bp. Based on the median NRLs and regularities inferred from these analyses, we named these clusters irregular-short (IRS), irregular-long (IRL), irregular-170 (IR170), regular repeat length 172 (NRL172), regular repeat length 187A and B (NRL187A/B), and regular repeat length 192 (NRL192). The difference between irregular and regular clusters is clear when closely inspecting histograms of NRL calls from selected clusters (**Figure 3D; Supplementary Figure 7C**), as well as the modification patterns on individual molecules (**Figure 3E**). Our analyses also varied with respect to the fraction of molecules per cluster where a secondary peak could be detected (0.50% - 38.2% of molecules across specific clusters; **Supplementary Figure 7D**). Failure to detect a peak within a single-molecule autocorrelogram could be due to multiple factors, including technical biases (*e.g*. random undermethylated molecules). We observed, however, that more ‘missing’ NRL estimates occurred in irregular clusters, suggesting that at least a fraction of failed peak calls occurred due to lack of intrinsic regularity along individual footprinted molecules. These analyses together demonstrate that SAMOSA data can be clustered in an unbiased manner, thus enabling estimates of the equilibrium composition of the genome with respect to oligonucleosome regularity and repeat length.

### SAMOSA captures the transient nucleosome occupancy of transcription factor binding motifs

We next explored the extent to which our data captures chromatin structure at predicted K562 transcription factor (TF) binding sites^37^. Both endo- and exo-nucleolytic MNase cleavage activities are obstructed by genomic protein-DNA contacts; resulting fragment-ends thus capture both nucleosomal- and TF-DNA interactions^38,39^. Inspection of cleavage patterns about 6 different transcription factor binding sites (CTCF, NRF1, NRSF / REST, PU.1, c-MYC, GATA1) (**Figure 4A-F**) revealed signal resembling traditional MNase-seq data, with fragment ends accumulating immediately proximal to predicted transcription factor binding motifs, and, in the case of some TFs (*i.e*. CTCF, REST, PU.1), showed characteristic patterns of phased nucleosomes. Further analysis of m^6^dA signal in sequenced molecules harboring motifs with at least 500 nucleotides of flanking DNA revealed examples of methyltransferase accessibility coincident with TF motifs (*e.g*. CTCF, NRF1, c-MYC), but also cases where single-molecule averages demonstrated weak or no differential signal when compared to equal numbers of molecules drawn from random genomic regions matched for GC-percentage and repeat content (*e.g*. GATA1; **Figure 4G-L**). Importantly, our methylation data do not appear to capture TF ‘footprints’ as seen in DNase I, hydroxyl radical, or MNase cleavage data—this could be due to turnover of transcription factors during our solubilization process, or owed to sterics, as EcoGII and other bacterial methyltransferases are roughly twice the molecular weight of *S. aureus* micrococcal nuclease^26^.

**Figure 4:**
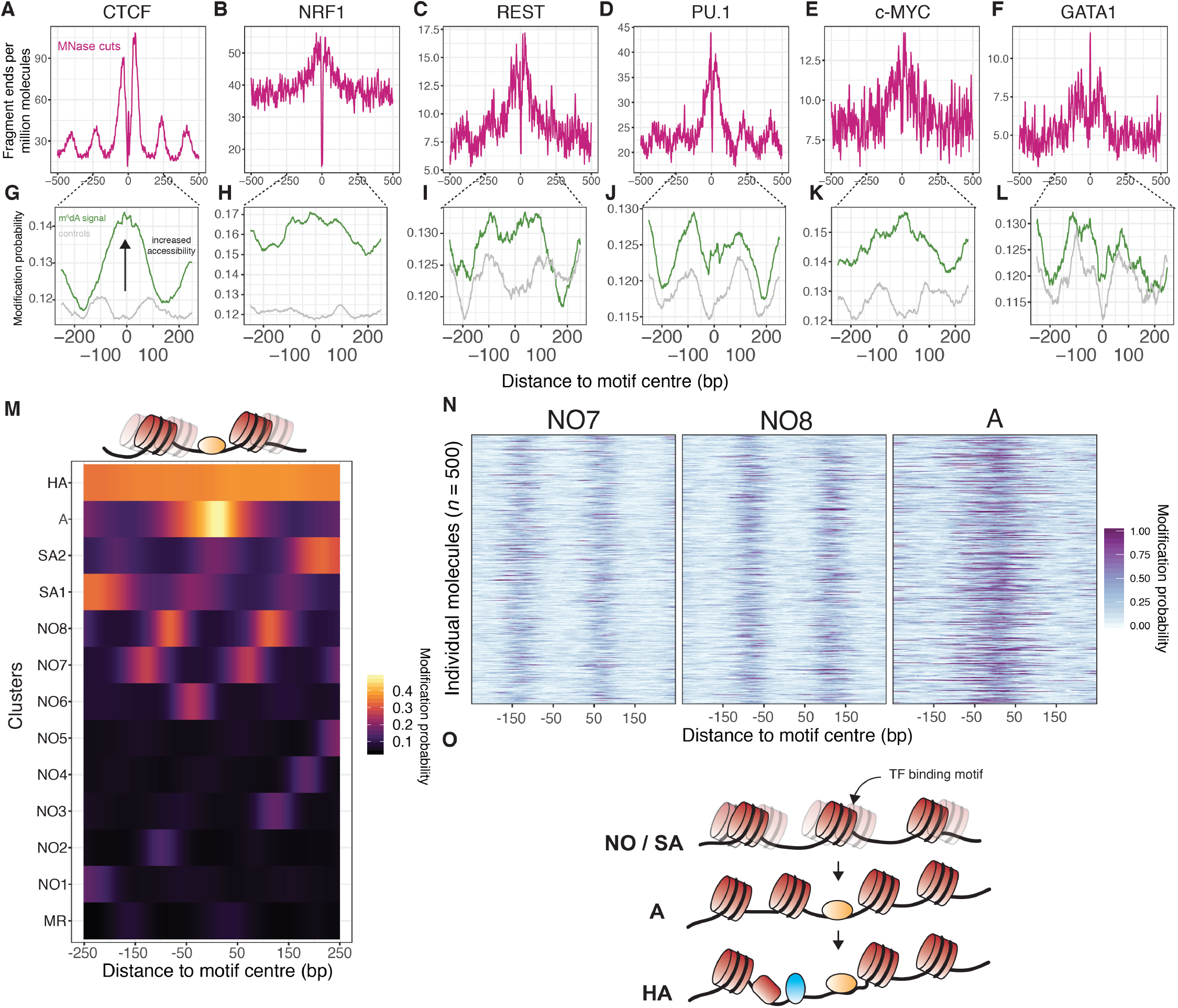
SAMOSA captures bulk and single-molecule evidence of transcription factor-DNA interaction simultaneously via two orthogonal molecular signals. **A.-F.)** SAMOSA MNase-cut signal averaged over predicted CTCF, NRF1, REST, PU.1, c-MYC, and GATA1 binding motifs in the K562 epigenome. All binding sites were predicted from ENCODE ChIP-seq data. **G-L.)** m^6^dA signal for the same transcription factors, averaged over molecules containing predicted binding sites and at least 250 bases flanking DNA on either side of the predicted motif. Methylation patterns at predicted sites were compared against average profiles taken from randomly drawn molecules from GC%- and repeat-content-matched regions of the genome (calculated for each ENCODE ChIP-seq peak set). **M.)** Results of clustering motif-containing molecules using the Leiden community detection algorithm. Clusters were manually annotated as containing molecules that were: ‘methylation resistant’ (MR), nucleosome occupied (NO1-8), stochastically accessible (SA1-2), accessible (A), or hyper-accessible (HA). **N.)** Heatmap representation of single-molecule accessibility profiles for clusters NO7, NO8, and A (500 randomly sampled molecules per cluster). **O.)** Our data may be explained by the Widom ‘site exposure’ model *in vivo*. Transcription factor binding motifs are stochastically exposed as nucleosomes toggle between multiple ‘registers’ as seen in **Figure 3M** (states NO and SA). Transcription factor binding perhaps enforces a favorable nucleosome register (state A), which can then seed hyper-accessible states / further TF-DNA interactions (state HA).

In theory, single-molecule footprinting data should distinguish nucleosome-bound and nucleosome-free states for molecules containing TF binding sites. These accessibility patterns should be specific to transcription factor binding motifs (*i.e*. not present in control molecules matched for GC / repeat content). To test whether our assay captured such signal, we clustered all molecules shown in **Figures 4G-L** (including control molecules) using Leiden clustering, using modification probabilities extracted in a 500 bp window surrounding the predicted motif site / control site. In total we defined thirteen discrete states of template accessibility across all surveyed molecules (**Figure 4M**; cluster sizes shown in **Supplementary Figure 8**). We interpreted these states on the basis of methyltransferase accessibility as: methyltransferase resistant motifs (MR); nucleosome-occluded motifs (NO1-8); stochastically-accessible motifs (wherein motif accessibility is slightly elevated near the DNA entry / exit point of a footprinted nucleosome; SA1-2); accessible motifs (A); and hyper-accessible motifs (HA). Notably, the patterns within these clusters were evident at single-molecule resolution (**Figure 4N**). We speculate that the broad distribution of these states across both TF binding sites and controls represent distributions of nucleosome ‘registers’ surrounding typical transcription factor binding motifs (*i.e*. states MR; NO-1-8). A fraction of these registers (*i.e*. states SA1/2) may stochastically permit transcription factor binding (perhaps through transient unwrapping of the nucleosome^40^), enabling formation of a new nucleosome register (*i.e*. state ‘A’), and subsequent generation of a highly accessible state (‘HA’; model illustrated in **Figure 4O**). The relative fraction of molecules in an ‘SA’ state could conceivably be modulated by TF intrinsic properties (*e.g*. ability to bind partially nucleosome-wrapped DNA^41^), or extrinsic factors (*e.g*. local concentration of ATP-dependent chromatin remodeling enzymes^42^). Indeed, most transcription factors (excepting PU.1 and GATA1—the latter of which may productively bind nucleosomal DNA^43^) were significantly enriched for specific states as defined above, and all control regions were markedly depleted for molecules harboring the accessible ‘A’ and ‘HA’ states, hinting at the biological relevance of these patterns (**Supplementary Figure 9**). Still, future experimental follow-up coupling our protocol with perturbed biological systems and deeper sequencing are necessary to quantitatively interrogate this model.

### Heterogeneous oligonucleosome patterns comprise human epigenomic domains

Short-read and long-read sequencing of nucleolytic fragments in mammals have suggested that NRLs vary across epigenomic domains, with euchromatin harboring shorter NRLs on average and heterochromatic domains harboring longer NRLs^44–46^, but the relative heterogeneity of these domains remains unknown. We speculated that SAMOSA data could be used to estimate single-molecule oligonucleosome pattern heterogeneity across epigenomic domains. We revisited the seven oligonucleosome patterns defined above, and examined the distribution of patterns across collections of single molecules falling within ENCODE-defined H3K4me3, H3K4me1, H3K36me3, H3K27me3, and H3K9me3-decorated chromatin domains. To control for the impact of GC-content on these analyses, we also included GC- / repeat-content matched control molecules for each epigenomic mark surveyed. Furthermore, to take advantage of the long-read and relatively unbiased nature of our data, we also incorporated molecules deriving from typically unmappable human alpha, beta, and gamma satellite DNA sampled directly from raw CCS reads.

We visualized the relative heterogeneity of these domains and controls in two ways: using histograms of computed single-molecule NRL estimates (**Figure 5A**), and by using stacked bar graphs to visualize cluster membership (**Figure 5B**). A striking finding of our analyses was that each epigenomic domain surveyed was comprised of a highly heterogeneous mixture of oligonucleosome patterns. In most cases, these patterns differed only subtly from control molecules with respect to regularity and NRL. In specific cases, we observed small effect shifts in the estimated median NRLs for specific domains—for example, a shift of ~5 bp (180 bp vs. 185 bp) in H3K9me3 chromatin with respect to random molecules, and a shift of ~4 bp (182 bp vs 186 bp) for H3K36me3. These shifts were also evident in the fraction of molecules with successful peak calls: H3K4me3 decorated chromatin, for example, had markedly fewer (78.0% vs 88.6%) successful calls compared to control molecules, a finding consistent with the expected irregularity of active promoter oligonucleosomes. We note that all of these measured parameters would be unattainable using any existing biochemical method, and that these preliminary findings argue against the abundance of homogeneous oligonucleosome structures in either heterochromatic or euchromatic nuclear regions.

**Figure 3:**
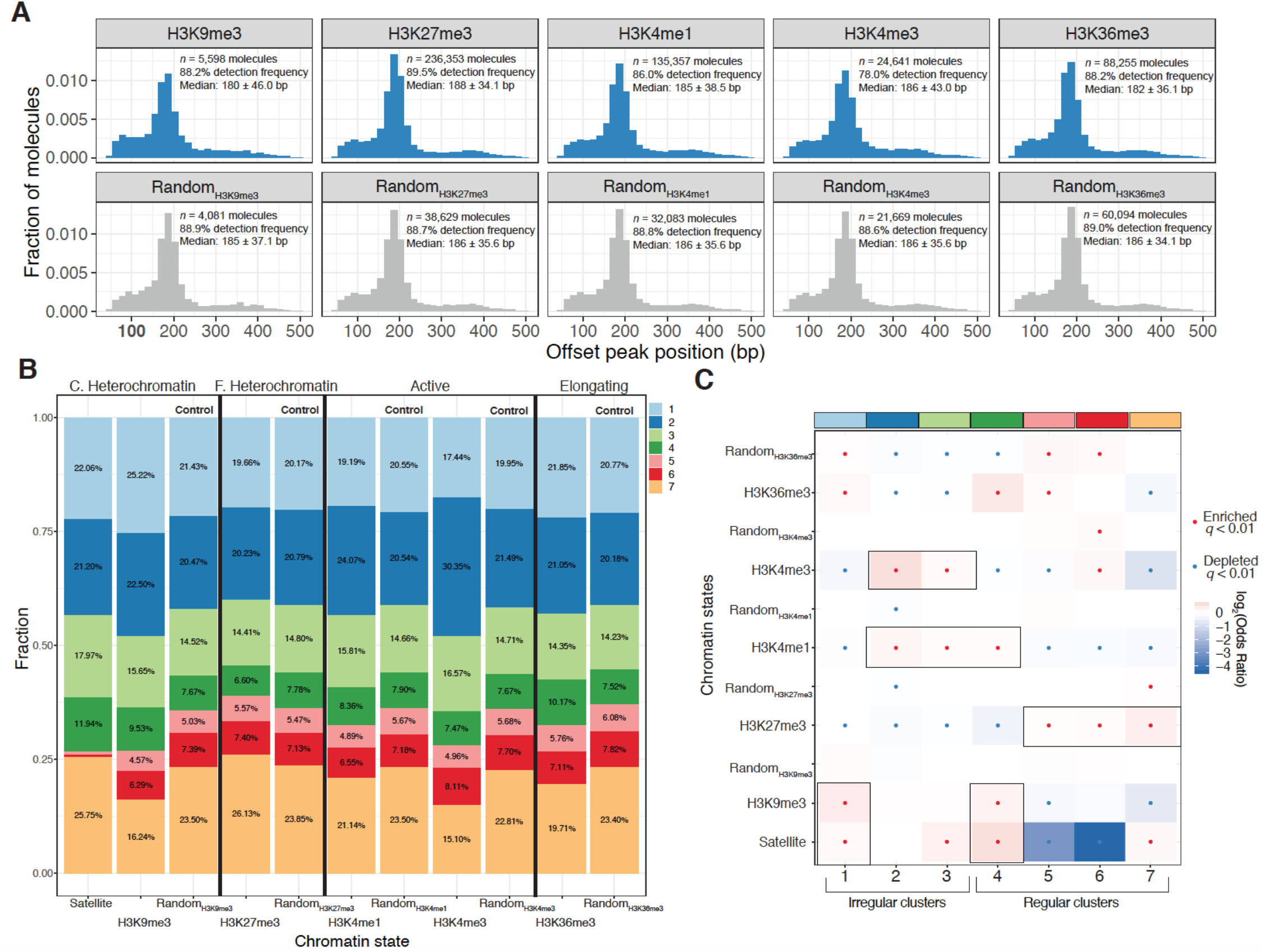
Human epigenomic states are punctuated by specific oligionucleosome patterns. **A.)** Histogram representations of the estimated single-molecule NRLs for five different epigenomic domains compared to control sets of molecules matched for GC and repeat content. *Inset*: Numbers of molecules plotted, median NRL estimates with associated median absolute deviations, and the percent of molecules where a peak could not be detected. **B.)** Stacked bar chart representation of the relative composition of each epigenomic domain with respect to the seven clusters defined in **Figure 3**. C. Heterochromatin: constitutive heterochromatin; F. Heterochromatin: facultative heterochromatin. **C.)** Heatmap of enrichment test results to determine nucleosome conformers that are enriched or depleted for each chromatin state. Tests qualitatively appearing to be chromatin-state specific are highlighted with a black box. Significant tests following multiple hypothesis correction marked with a black dot. Fisher’s Exact Test was used for all comparisons.

On first glance, our data appear to run counter to previous observations demonstrating that epigenomic domains can be delineated by differences in bulk nucleosome positioning as measured by nuclease digestion. One possible explanation for this is that epigenomic domains subtly, but significantly, vary in their relative composition of distinct oligonucleosome patterns, and the resulting average of these differences is the signal captured in MNase-Southern and other cleavage-based measurements. We tested this hypothesis by constructing a series of statistical tests to determine whether each of the seven defined oligonucleosome patterns were significantly enriched or depleted across chromatin domains and matched control regions (**Figure 5D**). Our results suggest that chromatin domains are demarcated by their relative usage of specific oligonucleosome patterns. Consistent with expectations, active chromatin marked by H3K4me3 and H3K4me1 are punctuated by a mixture of irregular oligonucleosome patterns (namely, clusters IRL and IR170). For transcription elongation associated H3K36me3 decorated chromatin, both short-read mapping in human and long-read bulk array regularity mapping in *D. melanogaster* have both suggested relatively short, regular nucleosome repeat lengths^20,44^. Our data partially corroborate this finding in human K562 cells: H3K36me3-domains are punctuated by irregular IRS oligoncleosome patterns (Fisher’s Exact Odds Ratio [O.R.] = 1.13; *q* = 1.71E-50) and regular, short NRL172 patterns (O.R. = 1.39; *q* = 3.69E-170).

The more uniform coverage of our assay also allows us to assess compositional biases in heterochromatic domains. Short-read-based human studies and classical MNase mapping of constituve heterochromatin have suggested that H3K9me3-decorated chromatin harbor i.) long nucleosome repeat lengths on average, and ii.) are highly regular. These estimates are susceptible to artifacts, as heterochromatic nucleosomes are expected to be both strongly phased and weakly positioned. Our data partially disagree with prior estimates—across both H3K9me3 and Satellite molecules we observe enrichment for irregular IRS nucleosome conformers (Satellite O.R. = 1.13; *q* = 5.71E-11; H3K9me3 O.R. = 1.35; *q* = 3.95E-23). Still, these enriched conformers were accompanied by enrichment for regular NRL172 oligonucleosome patterns for both states (Satellite O.R. = 1.61; *q* = 5.25E-80; H3K9me3 O.R. = 1.23; *q* = 3.86E-6). These analyses demonstrate that prior NRL estimates by short-read sequencing may have been confounded by *in vivo* heterogeneity in nucleosome positions, that heterochromatic nucleosome conformations can be both irregular and diverse, and finally, highlight the value of SAMOSA for accurately studying nucleosome structure in heterochromatin.

Taken as a whole, our data suggest two fundamental properties of human epigenomic domains: first, epigenomic domains are comprised of a diverse array of oligonucleosome patterns varying substantially in intrinsic regularity and average distance between regularly spaced nucleosomes; second: epigenomic domains are demarcated by their usage of these oligonucleosome patterns. We find that all epigenomic states are characterized by a diverse mixture of oligonucleosomal conformers—many conformational states are neither significantly depleted nor enriched with respect to all molecules surveyed, further hinting at the diverse composition of chromatin domains genome-wide.

## DISCUSSION

Here, we present the <u>S</u>ingle-molecule <u>A</u>denine <u>M</u>ethylated <u>O</u>ligonucleosome <u>S</u>equencing <u>A</u>ssay (SAMOSA), a method for resolving nucleosome-DNA interactions using the EcoGII adenine methyltransferase and PacBio single-molecule real-time sequencing. Our approach has multiple advantages over existing methyltransferase-based sequencing approaches: first, by using a relatively nonspecific methyltransferase, we avoid the primary sequence biases associated with GpC / CpG methyltransferase footprinting methods; second, by natively detecting modifications using the single-molecule real time sequencer, we reduce enzymatic sequence bias and avoid sample damage associated with sodium bisulphite conversion; third, by employing a light MNase-digest and solubilizing long oligonucleosomal fragments, we more uniformly capture both active and silent chromatin and simultaneously measure methyltransferase accessibility and MNase cleavage on single molecules; finally, and most importantly, our approach unlocks the study of protein-DNA interactions at length-scales previously unallowed by Illumina sequencing.

Our study does have limitations. While the current SAMOSA protocol enriches fragments ranging from ~500 bp - ~2 kb in size, high-quality PacBio CCS sequencing is compatible with fragments ranging from 10 – 15 kbp. We anticipate that with further optimization, SAMOSA will be applicable to longer arrays, enabling kilobase-domain-scale study of single-molecule oligonucleosome patterning. Our approach also requires solubilization of oligonucleosomal fragments, and is thus unlikely to capture protein-DNA interactions weaker or more transient than the stable nucleosome-DNA interaction. Third, our approach does not bias signal towards regions of open chromatin (as in short-read DNase / ATAC-seq or long-read SMAC-seq^23^), and thus addresses a different set of gene regulatory questions compared to those published assays. Fourth, our proof-of-concept was performed in unsynchronized K562 cells, and thus we cannot yet addres the contribution of a biological process like the cell cycle to the observed heterogeneity. Finally, as a proof-of-concept our approach falls short of generating a high-coverage reference map of the K562 epigenome; as sequencing costs for PacBio decrease and sequence-enrichment technologies (*e.g*. CRISPR-based enrichment^47^; SMRT-ChIP^48^) for the platform mature, SAMOSA may routinely be used to generate reference datasets with hundred-to-thousand-fold single-molecular coverage of genomic sites of interest.

Our data confirms that the human epigenome is made up of a diverse array of oligonucleosome patterns, including highly regular arrays of varying nucleosome repeat lengths, and irregular arrays where nucleosomes are positioned without a detectable periodic signature^49^. Our results broadly agree with a recent approach employing electron tomography to map the *in situ* structure of mammalian nuclei, which found chromatin to be highly heterogeneous at the length scale of multi-nucleosome interactions, and failed to detect evidence of a 30 nm fibre or other homogeneous higher-order compaction states^7^. At the sequencing depth presented here, these oligonucleosome patterns significantly, if subtly, vary across different epigenomic domains. Surprisingly, we find that both mappable (H3K9me3 ChIP-seq peaks) and unmappable (human satellite sequence) constitutive heterochromatin are enriched for irregular oligonucleosome patterns in addition to expected regular arrays—the presence of these irregular fibres may have been previously missed due to an understandable reliance on bulk averaged methods (*e.g*. MNase-Southern) for studying constitutive heterochromatin. Indeed, it is tempting to speculate that this irregularity may be linked to the dynamic restructuring of heterochromatic nucleosomes by factors like HP1^50^, which may promote phase-separation of heterochromatin. Future studies combining SAMOSA with cellular perturbation of heterochromatin-associated factors could directly address this.

More generally, future work employing our technique must focus on questioning the biological significance of this global heterogeneity: for example, is the fraction of stochastically accessible transcription factor binding sites (*i.e*. motif ‘site exposure’ frequency^40,51^) important for TF-DNA binding in nucleosome-occluded genomic regions? What is the interplay between transcription factor ‘pioneering’ and stochastic site accessibility? What are the global roles of ATP-dependent chromatin remodeling enzymes (*i.e*. SWI/SNF; ISWI; INO80; CHD) in maintaining these patterns genome-wide^52^? Our approach also unlocks a set of conceptual questions regarding the nature of chromatin secondary structure. Significant genome-wide efforts have revealed that metazoan epigenomes are punctuated by regions of concerted histone modification and subnuclear positioning^37,53^, but approaches for studying the distribution of oligonucleosomal patterns associated within these same regions are lacking. Given recent work suggesting that NRLs can specify the ability of nucleosomal arrays to phase separate^54^, it is likely that SAMOSA and similar assays may provide an important bridge between *in vitro* biochemical observations of chromatin and *in vivo* genome-wide ‘catalogs’ of oligonucleosome patterning.

SAMOSA adds to the growing list of technologies that use high-throughput single-molecule sequencing to explore the epigenome^20,23,24,31^. We foresee the broad applicability of this and similar approaches to dissect gene regulatory processes at previously intractable length-scales. Our approach and associated analytical pipelines demonstrate the versatility of high-throughput single-molecule sequencing—namely the ability to cluster single-molecules in an unsupervised manner to uncover molecular states previously missed by short-read approaches. Our analytical approach bears many similarities to methods used in single-cell analysis, and indeed many of the technologies and concepts typically used for single-cell genomics^55^ (*e.g*. clustering; trajectory analysis) will likely have value when applied to single-molecule epigenomic assays. Our approach also represents the first of what we anticipate will be many ‘multi-omic’ single-molecule assays; as third-generation sequencing technologies advance, it will likely become possible to encode multiple biochemical signals on the same single-molecules, thus enabling causal inference of the logic and ordering of biochemical modifications on single chromatin templates.

## METHODS

### Preparation of nonanucleosome arrays via Salt Gradient Dialysis

The nonanucleosome DNA in a plasmid was purified by Gigaprep (Qiagen) and the insert was digested out with EcoRV, ApaLI, XhoI and StuI. The insert was subsequently purified using a Sephacryl S1000 super fine gel flitration (GE Healthcare). Histones were purified and octamer was assembled as previously described^56^. To assemble the arrays, the nonanucleosome DNA was mixed with octamer and supplementing dimer, then dialyzed from high salt to low salt^57^. EcoRI sites engineered in the linker DNA between the nucleosomes, and digestion by EcoRI was used to assess the quality of nucleosome assembly.

### SAMOSA on nonanucleosomal chromatin arrays

For the chromatin arrays, 1.5 ug of assembled array was utilized as input for methylation reactions with the non-specific adenine EcoGII methyltransferase (New England Biolabs, high concentration stock; 2.5E4U / mL). For the naked DNA controls, 2 ug of DNA was utilized as input for methylation reactions. Methylation reactions were performed in a 100 uL reaction with Methylation Reaction buffer (1X CutSmart Buffer,1 mM S-adenosyl-methionine (SAM, New England Biolabs)) and incubated with 2.5 uL EcoGII at 37°C for 30 minutes. SAM was replenished to 6.25 mM after 15 minutes. Unmethylated controls were similarly supplemented with Methylation Reaction buffer, minus EcoGII and replenishing SAM, and the following purification conditions. To purify DNA, the samples were all subsequently incubated with 10uL Proteinase K (20mg/mL) and 10uL 10% SDS at 65°C for a minimum of 2 hours up to overnight. To extract the DNA, equal parts volume of Phenol-Chloroform was added and mixed vigorously by shaking, spun (max speed, 2 min). The aqueous portion was carefully removed and 0.1x volumes of 3M NaOAc, 3uL of GlycoBlue and 3x volumes of 100% EtOH were added, mixed gently by inversion, and incubated overnight at −20°C. Samples were then spun (max speed, 4°C, 30 min), washed with 500uL 70% EtOH, air dried and resuspended in 50uL EB. Sample concentration was measured by Qubit High Sensitivity DNA Assay.

### Preparation of *in vitro* SAMOSA SMRT Libraries

The purified DNA from nonanucleosome array and DNA samples were used in entirety as input for PacBio SMRTbell^®^ library preparation (~1.5-2 ug). Preparation of libraries included DNA damage repair, end repair, SMRTbell^®^ ligation, and Exonuclease according to manufacturer’s instruction. After Exonuclease Cleanup and a double 0.8x Ampure PB Cleanup, sample concentration was measured by Qubit High Sensitivity DNA Assay (1 uL each). To assess for library quality, samples (1 uL each) were run on an Agilent Bioanalyzer DNA chip. Libraries were sequenced on either Sequel I or Sequel II flow cells (UC Berkeley QB3 Genomics). Sequel II runs were performed using v2.0 sequencing chemistry and 30 hour movies.

### Cell lines and cell culture

K562 cells (ATCC) were grown in standard media containing RPMI 1640 (Gibco) supplemented with 10% Fetal Bovine Serum (Gemini, Lot#A98G00K) and 1% Penicillin-Streptomycin (Gibco). Cell lines were regularly tested for mycoplasma contamination and confirmed negative with PCR (NEB NebNext Q5 High Fidelity 2X Master Mix).

### Isolation of nuclei, MNase digest, and overnight dialysis

100E6 K562 cells were collected by centrifugation (300x**g**, 5 min), washed in ice cold 1X PBS, and resuspended in 1 mL Nuclear Isolation Buffer (20mM HEPES, 10mM KCl, 1mM MgCl2, 0.1% Triton X-100, 20% Glycerol, and 1X Protease Inhibitor (Roche)) per 5-10 e6 cells by gently pipetting 5x with a wide-bore tip to release nuclei. The suspension was incubated on ice for 5 minutes, and nuclei were pelleted (600xg, 4°C, 5 min), washed with Buffer M (15mM Tris-HCl pH 8.0, 15 mM NaCl, 60mM KCl, 0.5mM Spermidine), and spun once again. Nuclei were resuspended in 37°C pre-warmed Buffer M supplemented with 1mM CaCl_2_ and distributed into two 1mL aliquots. For digestion, micrococcal nuclease from *Staphylococcus aureus* (Sigma, reconstituted in ddH2O, stock at 0.2U/uL) was added at 1U per 50E6 nuclei, and nuclei were digested for 1 min. at 37°C. EGTA was added to 2mM immediately after 1 minute to stop the digestion and incubated on ice. For nuclear lysis and liberation of chromatin fibres, MNase-digested nuclei were collected (600xg, 4°C, 5 minutes) and resuspended in 1mL per 50E6 nuclei of Tep20 Buffer (10mM Tris-HCl pH 7.5, 0.1mM EGTA, 20mM NaCl, and 1X Protease Inhibitor (Roche) added immediately before use) supplemented with 300ug/mL of Lysolethicin (L-α-Lysophosphatidylcholine from bovine brain, Sigma, stock at 5mg/mL) and incubated at 4°C overnight. To remove nuclear debris the next day, dialyzed samples were spun (12,000xg, 4°C, 5 minutes) and the soluble chromatin fibres present in the supernatant were collected. Sample concentration was measured by Nanodrop.

### SAMOSA on K562-derived oligonucleosomes

Dialyzed chromatin was utilized as input (1.5 ug) for methylation reactions with the non-specific adenine EcoGII methyltransferase (New England Biolabs, high concentration stock 2.5e4U/mL). Reactions were performed in a 200uL reaction with 1X CutSmart Buffer and 1mM S-adenosyl-methionine (SAM, New England Biolabs) and incubated with 2.5uL enzyme at 37°C for 30 minutes. SAM was replenished to 6.25mM after 15 minutes. Non-methylation controls were similarly supplemented with Methylation Reaction buffer, minus EcoGII and replenishing SAM, and purified by the following conditions. To purify all DNA samples, reactions were incubated with 10uL of RNaseA at room temperature for 10 minutes, followed by 20uL Proteinase K (20mg/mL) and 20uL 10% SDS at 65°C for a minimum of 2 hours up to overnight. To extract the DNA, equal parts volume of Phenol-Chloroform was added and mixed vigorously by shaking, spun (max speed, 2 min). The aqueous portion was carefully removed and 0.1x volumes of 3M NaOAc, 3uL of GlycoBlue and 3x volumes of 100% EtOH were added, mixed gently by inversion, and incubated overnight at −20°C. Samples were then spun (max speed, 4°C, 30 min), washed with 500uL 70% EtOH, air dried and resuspended in 50uL EB. Sample concentration was measured by Qubit High Sensitivity DNA Assay. Naked DNA Positive methylation controls were collected from aforementioned non-methylated controls post-purification (25uL, ~500ng), methylated with EcoGII as previously stated, and purified again by the following conditions.

### Preparation of *in vivo* SAMOSA SMRT libraries

Purified DNA from MNase-digested K562 chromatin oligonucleosomes (methylated, non-methylated control, purified then methylated) were briefly spun in a Covaris G-Tube (3380xg, 1 min) in efforts to shear gDNA uniformly to 10kB prior PacBio library preparation. The input concentration was approximately 575ng for methylated and non-methylated samples, and approximately 320ng for purified then methylated samples. Samples were concentrated with 0.45x of AMPure PB beads according to manufacturer’s instructions. The entire sample volume was utilized as input for subsequent steps in library preparation, which included DNA damage repair, end repair, SMRTbell ligation, and Exonuclease cleanup according to manufacturer’s instructions. For SMRTbell ligations, unique PacBio SMRT-bell adaptors (100uM stock) were annealed to a 20uM working stock in 10mM Tris-HCl pH 7.5 and 100mM NaCl in a thermocycler (85°C 5 min, RT 30sec, 4°C hold) and stored at −20°C for long-term storage. After exonuclease cleanup and double Ampure PB cleanups (0.45X), the sample concentrations were measured by Qubit High Sensitivity DNA Assay (1uL each). To assess for size distribution and library quality, samples (1 uL each) were run on an Agilent Bioanalyzer DNA chip. Libraries were sequenced on Sequel II flow cells (UC Berkeley QB3 Genomics Core). *In vivo* data were collected over two 30 h Sequel II movie runs; the first with a 2 h pre-extension time and the second with a 0.7 h pre-extension time.

### Data analysis

All raw data will be made available at GEO Accession XXXXXX; processed data is available at Zenodo (dx.doi.org/10.5281/zenodo.3834706). All scripts and notebooks for reproducing analyses in the paper are available at https://github.com/RamaniLab/SAMOSA.

We apply our method to two use cases in the paper, and they differ in the computational workflow to analyze them. The first is for sequencing samples where every DNA molecule should have the same sequence, which is the case for our *in vitro* validation experiments presented in Figure 1. The second use case is for samples from cells containing varied sequences of DNA molecules. We will refer to the first as homogeneous samples, and the second as genomic samples. The workflow for genomic samples will be presented first in each sections, and the deviations for homogeneous samples detailed at the end.

#### Sequencing read processing

Sequencing reads were processed using software from Pacific Biosciences. The following describes the workflow for genomic samples:

1. **Demultiplex reads** Reads were demultiplexed using lima. The flag ‘--same’ was passed as libraries were generated with the same barcode on both ends. This produces a BAM file for the subreads of each sample.
2. **Generate Circular Consensus Sequences (CCS)** CCS were generated for each sample using ccs^58^. Default parameters were used other than setting the number of threads with ‘-j’. This produces a BAM file of CCS.
3. **Align CCS to the reference genome** Alignment was done using pbmm2^59^, and run on each CCS file, resulting in BAM files containing the CCS and alignment information.
4. **Generate missing indices** Our analysis code requires pacbio index files (.pbi) for each BAM file. ‘pbmm2’ does not generate index files, so missing indices were generated using ‘pbindex’. For homogeneous samples, replace step 3 with this alternate step 3
5. **Align subreads to the reference genome** pbmm2 was run on each subreads BAM file (the output of step 1) to align subreads to the reference sequence, producing a BAM file of aligned subreads.

#### Sample Reference Preparation

Our script for analyzing samples relies on a CSV file input that contains information about each sample, including the locations of the relevant BAM files and a path to the reference genome. The CSV needs a header with the following columns:

**index**: Integer indices for each sample. We write the table using ‘pandas’ ‘.to_csv’ function, with parameters ‘index=True, index_label-‘index’’
**cell**: A unique name for the SMRT cell on which the sample was sequenced **sampleName**: The name of the sample
**unalignedSubreadsFile**: This will be the file produced by step 1 above. This should be an absolute path to the file.
**ccsFile**: This is the file produced by step 2 above **alignedSubreadsFile**: This is the file produced by the alternate step 3 above. It is required for homogeneous samples but can be left blank for genomic samples.
**alignedCcsFile**: This is the file produced by step 3 above. It is required for genomic samples but can be left blank for homogeneous samples.
**reference**: The file of the reference genome or reference sequence for the sample

#### Extracting IPD measurements and calling methylation

The script extractIPD.py accesses the BAM files, reads the IPD values at each base and uses a gaussian mixture model to generate posterior probabilities of each adenine being methylated. extractIPD takes two positional arguments. The first is a path to the above sample reference CSV file. The second is a specification for which sample to run on. This can be either an integer index value, in which case extractIPD will run on the corresponding row. Alternatively it can be a string containing the cell and sampleName, separated by a period. Either way extractIPD will run on the specified sample using the paths to the BAM files contained within the CSV.

extractIPD produces the following three output files when run on genomic samples:

**processed/onlyT/{cell}_{sampleName}_onlyT_zmwinfo.pickle:** This file is a ‘pandas’ dataframe stored as a pickle, and can be read with the ‘pandas.read_pickle’ function. This dataframe contains various information about each individual ZMW.
**processed/onlyT/{cell}_{sampleName}_onlyT.pickle**: This file contains the normalized IPD value at every thymine. The data is stored as a dictionary object. The keys are the ZMW hole numbers (stored in the column zmw in the zmwinfo dataframe), and the values are numpy arrays. The arrays are 1D with length equal to the length of the CCS for that molecule. At bases that are A/T, there will be a normalized IPD value. Each G/C base and a few A/T bases for which an IPD value couldn’t be measured will contain NaN.
**processed/binarized/{cell}_{sampleName}_bingmm.pickle**: This file contains the posterior probability of each adenine being methylated. The data format is identical to the _onlyT.pickle file above, except the numpy array contains values between 0 and 1, where the higher values indicate a higher confidence that the adenine is methylated.

When run on homogeneous samples the following output files are alternately produced:

**processed/onlyT/{cell}_{sampleName}_onlyT.npy:** This numpy array has a column for every base in the reference sequence, and a row for each DNA molecule that passes the filtering threshold. A normalized IPD value is stored for each adenine that could be measured at A/T bases, other bases are NaN.
**processed/binarized/{cell}_{sampleName}_bingmm.npy:** This numpy array is the same shape as the _onlyT.npy file above. The values are posterior probabilities for an adenine being methylated, ranging from 0 to 1.

#### Dyad calling on in vitro methylated chromatin arrays

Nucleosome positions were predicted in nonanucleosomal array data by taking a 133 bp wide rolling mean across the molecule, and finding each local minimum peak at least 147 bp apart from each other.

#### In vivo *analyses*

We smooth the posterior probabilities calculated in the paper to account for regions with low local A/T content and generally denoise the single-molecule signal. For *in vitro* analyses, we smooth the calculated posterior probabilities using a 5 bp rolling mean. For all *in vivo* analyses in the paper that involve calculation of single-molecule autocorrelograms, averaging over multiple templates, and visualizing individual molecules, we smooth posteriors with a 33 bp rolling mean. For all autocorrelation calculations we ignore regions where compared lengths would be unequal; this has the effect of rendering the returned autocorrelogram exactly 0.5 * the input length.

#### Averages of the modification signal across the first 1 kb of K562 oligonucleosomes

We took all molecules at least 500 nt in length and concatenated all of the resulting matrices from each of the four separate samples / runs, and then plotted the NaN-sensitive mean over the matrix as a function of distance along the molecule.

#### ChromHMM Coverage Enrichment Analysis

We downloaded K562 ChromHMM labels from the UCSC Genome Browser, lifted over coordinates to hg38 using the liftOver tool, and then used bedtools multicov to compute the read coverage for each ChromHMM BED entry. We then read this file in as a pandas dataframe. We next use this dataframe to estimate the relative enrichment / depletion of each label in each dataset. The datasets we use in addition to our in-house SAMOSA data, we use aligned ENCODE^37^ DNaseI-seq data (ENCFF156LGK), in-house aligned K562 MNase-seq data^34^ and publicly-released whole-genome shotgun sequencing CCS data from PacBio (https://downloads.pacbcloud.com/public/dataset/HG002_SV_and_SNV_CCS/consensusreads/). We estimated enrichment by evaluating normalizing the number of reads mapping to each ChromHMM labeled genomic bin, dividing this value by the total number of bins with that ChromHMM label, and taking the natural log. We then plotted this data in heatmap form using ggplot2.

#### Clustering analysis of all chromatin molecules >=500 bp in length

We used Leiden clustering cluster all molecules in our dataset passing our lower length cutoff. Resolution and n_neighbors were manually adjusted to avoid generating large numbers of very small clusters (i.e. < 100 molecules). All parameters used for plotting figures in the paper are recapitulated in the Jupyter notebook. Our clustering strategy was as follows: first, we smoothed raw signal matrices with a 33 bp NaN-sensitive running mean. We next computed the autocorrelation function for each molecule in the matrix, using the full length of the molecule up to 1000 bp. We then used Scanpy^60^ to perform Leiden clustering on the resulting matrix. We visualized the resulting cluster averages with respect to the average autocorrelation function, and with respect to averaged modification probabilities for each cluster. For a subset of clusters we also randomly sampled 500 - 5000 molecules to directly visualize in the paper.

#### Computing single-molecule autocorrelograms and estimating NRLs on single molecules

We computed single-molecule autocorrelograms and discovered peaks on these autocorrelograms as follows: for each molecule, we used the scipy^61^ find_peaks function to in the computed autocorrelogram and annotated the location of that peak. We also kept track of the molecules where find_peaks could not detect a peak using the given parameters, which we optimized manually by modifying peak height / width to detect peaks on the averaged autocorrelograms. In our hands, these parameters robustly detect peaks between 180-190 bp in auto-correlgram averages, consistent with the expected bulk NRL in K562 cells (analyses by A Rendeiro; zenodo.org/record/3820875). For each collection of single-molecule autocorrelogram peaks we computed the median, the median absolute deviation, and visualized the distribution of peak locations as a histogram.

#### Transcription factor binding motif analyses and enrichment tests

K562 transcription factor binding sites were predicted as in Ramani *et al*^39^. Briefly, we downloaded IDR-filtered ENCODE ChIP-seq peaks for CTCF, NRF1, REST, c-MYC, PU.1, and GATA1, and then used FIMO^62^ to predict TF binding sites within these peaks using CISTROME PWM definitions for each transcription factor. For MNase-cleavage analyses, we plotted the abundance of MNase cuts (2 per molecule) with respect to TF binding sites and plotted these as number of cleavages per molecules sequenced. To examine modification probabilities around TF binding sites, we wrote a custom script (zmw_selector.py) to find the ZMWs that overlap with features of interest (e.g. transcription factor binding sites). We extracted all ZMWs where a portion of the read alignment falls within 1 kb of a given feature, and annotated the position of the alignment starts, ends, and strand with respect to the feature. We then used these coordinates and strand information to extract all modification signal falling within a 500 bp window centered at each TF binding site. For control sites, we used the gkmSVM package^63^ to find GC- / repeat-content matched genomic regions for each peakset. We constructed a series of enrichment tests (Fisher’s Exact) to determine odds ratios / p-values to find specific cluster label--transcription factor pairs that were enriched with respect to the total set of all labeled molecules. Finally, we used the Storey q-value package^64^ to correct for the number of Fisher’s exact tests performed.

#### Enrichment Tests for Chromatin States

We used a custom python script (zmw_selector_bed.py) or directly scanned for satellite-containing CCS reads (see below) to extract molecules that fall within ENCODE-defined chromatin states / pertain to human major satellite sequences. We then used a Python dictionary linking ZMW IDs to indices along the total matrix of molecules to link Cluster IDs and chromatin states. Finally, we constructed a series of enrichment tests (Fisher’s Exact) to determine odds ratios / p-values to find specific cluster label-chromatin state pairs that were enriched with respect to the total set of all labeled molecules. We then used the Storey q-value package to correct for the number of Fisher’s exact tests performed. Control molecules were drawn as above, using the gkmSVM package to find GC / repeat-content matched genomic regions for each peakset.

#### Selection of satellite-containing reads

Circular consensus reads with minimum length of 1-kb bearing satellites were identified using BLAST searching against a database containing DFAM^65^ consensus sequences for alpha (DF0000014.4, DF0000015.4, DF0000029.4), beta (DF0000075.4, DF0000076.4, DF0000077.4, DF0000078.4, DF0000079.4), and gamma (DF0000148.4, DF0000150.4, DF0000152.4) satellites using blastn with default parameters. Satellite containing reads were further filtered such that they contained at minimum two hits to satellite consensus sequences and matches spanned at least 50% of the consensus sequence.

## AUTHOR CONTRIBUTIONS

N.J.A., L.J.H., A.K., V.R. performed experiments. C.P.M., S.K., L.E., M.K., V.R. performed analyses. J.G.U. aided with experimental design and PacBio instrument troubleshooting. H.G., G.J.N., V.R. supervised research. N.J.A., C.P.M., V.R. wrote the manuscript with input from all authors.

## ACKNOWLEDGEMENTS

The authors thank Daniele Canzio (UCSF), Hiten Madhani (UCSF), Srinivas Ramachandran (CU Denver), and Christopher Weber (Stanford) for helpful discussions and comments on the manuscript. The authors thank Shana McDevitt, Robert Munch, and the UC Berkeley Vincent J Coates Genomics Sequencing Laboratory for assisting with PacBio sequencing. L.J.H. is supported by an ACS postdoctoral fellowship awarded the Roaring Fork Valley Group. This work was funded by NIH grant R01GM123977 to H.G., NIH grant R35GM127020 to G.J.N., and through the gracious support of the Sandler Foundation and the UCSF Program for Breakthroughs in Biomedical Research (to V.R.).

## HIGH-RESOLUTION SUPPLEMENTARY FIGURES

**Supplementary Figure 1:**
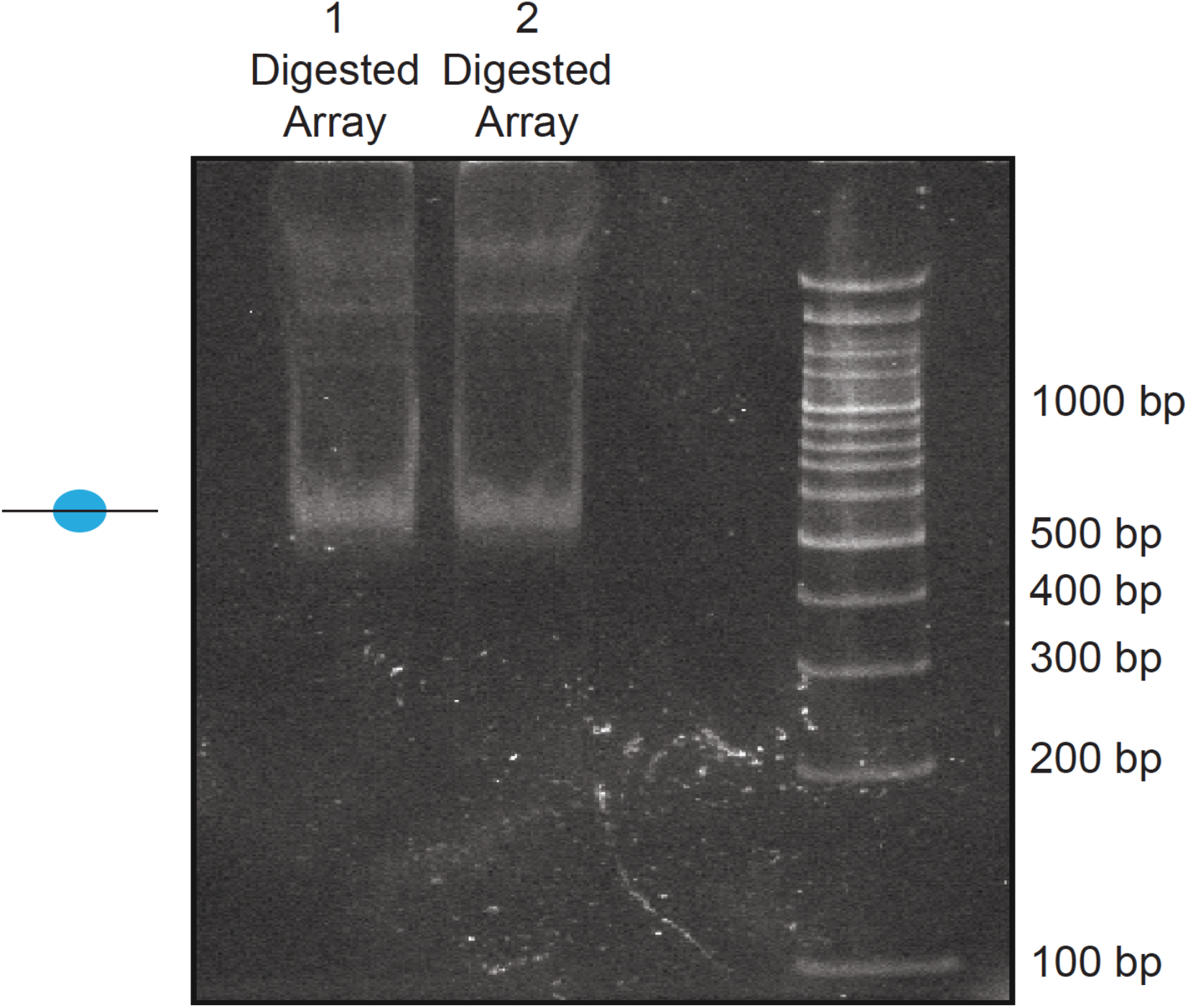
Quality control of *in vitro* nucleosome arrays assembled through salt-gradient dialysis. Nonanucleosomal arrays were assembled in duplicate as previously and checked for assembly extent via restriction enzyme digest. In both cases, the smallest digestion products corresponded to a protected mononucleosomal fragment, suggesting that there is minimal underassembly of the resulting arrays.

**Supplementary Figure 2:**
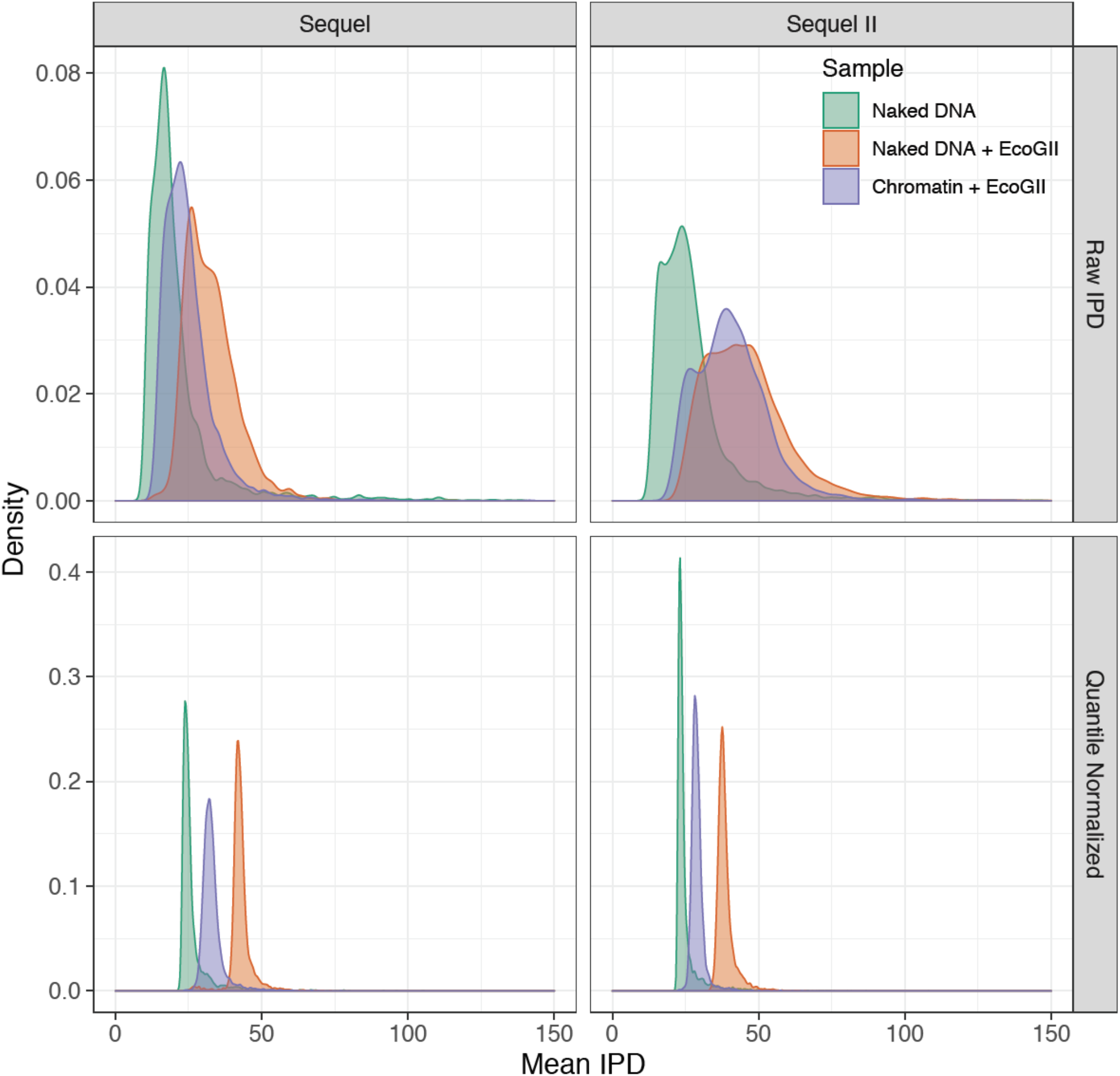
Mean raw and quantile normalized interpulse durations for *in vitro* SAMOSA experiments. *In vitro* SAMOSA experiments demonstrate intermediate single-molecule average interpulse durations compared to unmethylated array DNA and fully methylated deproteinated array DNA. Data are similarly separated on the Sequel and Sequel II platforms and quantile normalization further aids in separating chromatin from control samples, particularly on the Sequel II platform.

**Supplementary Figure 3:**
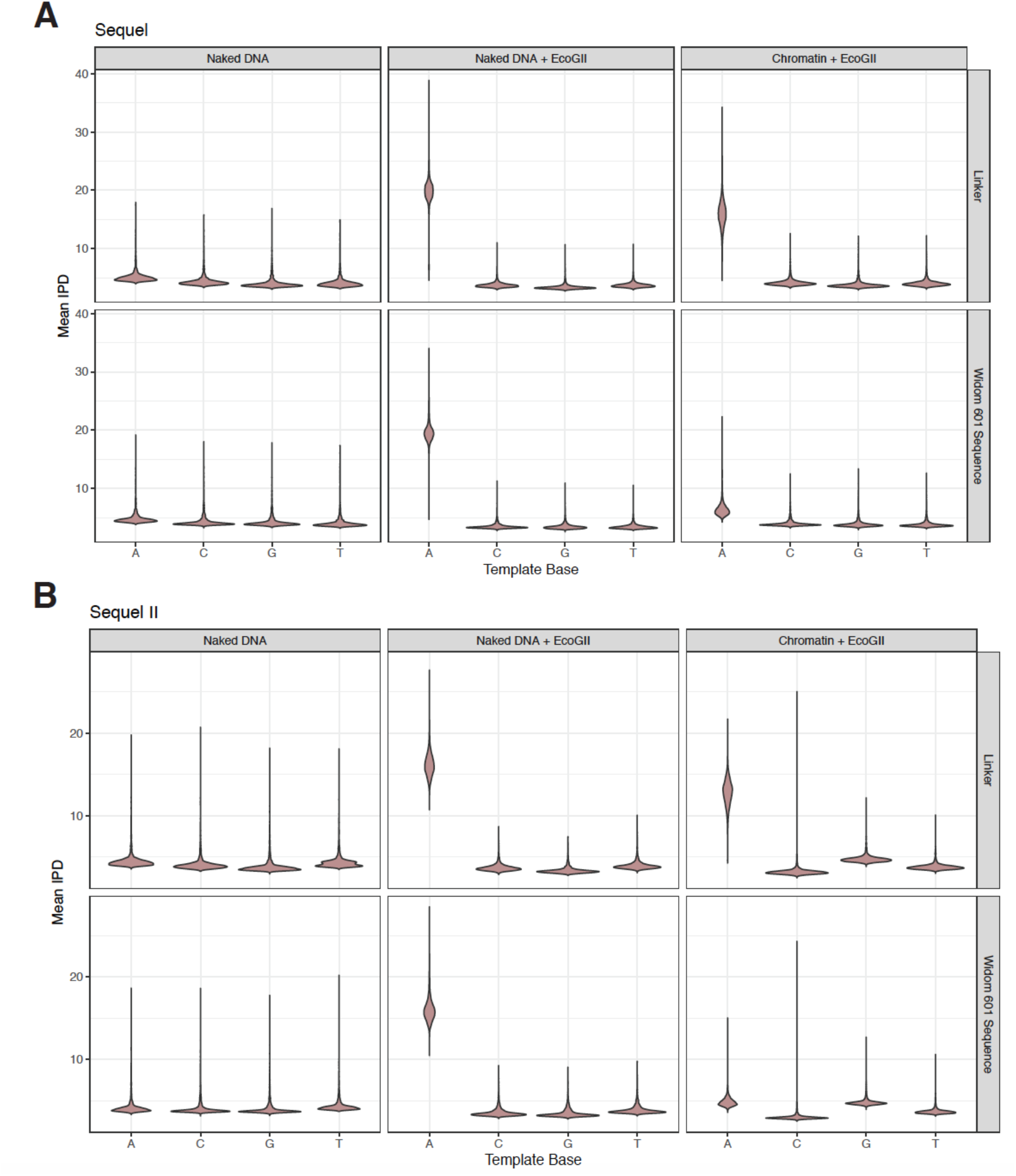
Adenine methylation by the EcoGII enzyme is specific to accessible adenines and is protected against by the nucleosome. **A.)** Violin plots of the distribution of average, quantile normalized IPD values for each nucleotide on controls (unmethylated / fully methylated naked DNA) and chromatin separated by nucleotides falling within the Widom 601 nucleosome positioning sequence or linker DNA. In the chromatin context, only adenine nucleotides falling within the linker are modified, consistent with protection of bases by nucleosomes positioned by the Widom 601 sequence. **B.)** As in A, but for data from the Sequel II platform.

**Supplementary Figure 4:**
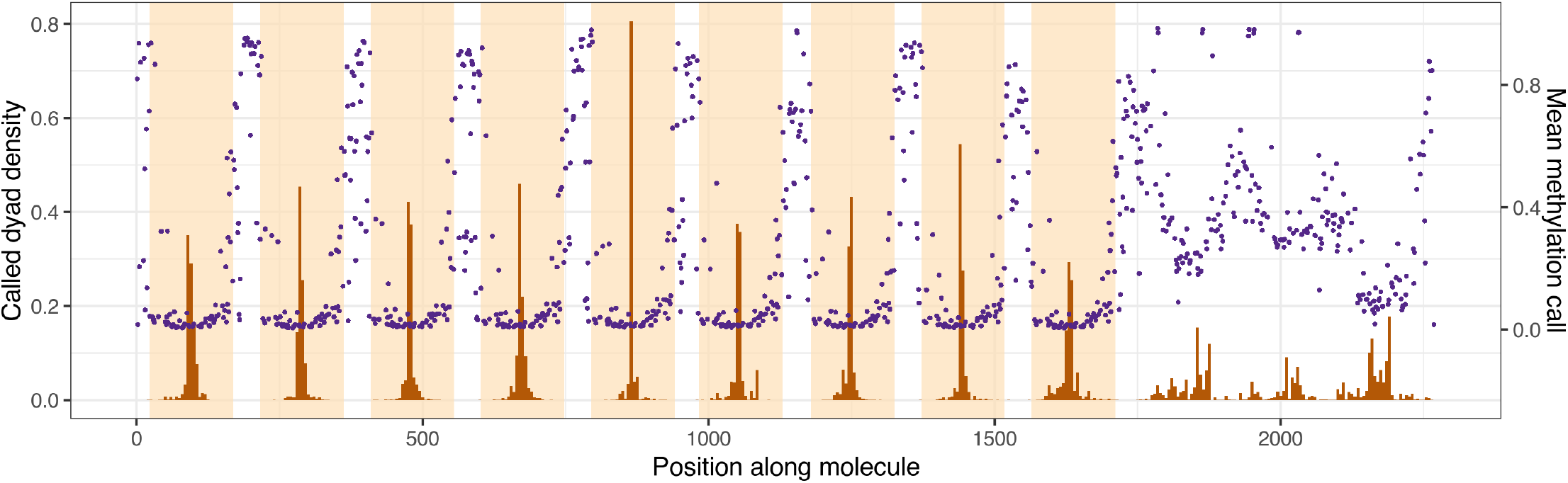
Average linker methylation and individually called dyad positions are qualitatively similar across the length of the nonanucleosomal array molecule. Histograms of called dyad positions for each occurrence of a Widom 601 repeat unit (orange shading), averaged over all sequenced chromatin molecules are shown in brown. Mean methylation calls for each linker sequence (sequence outside orange shading) are shown in purple.

**Supplementary Figure 5:**
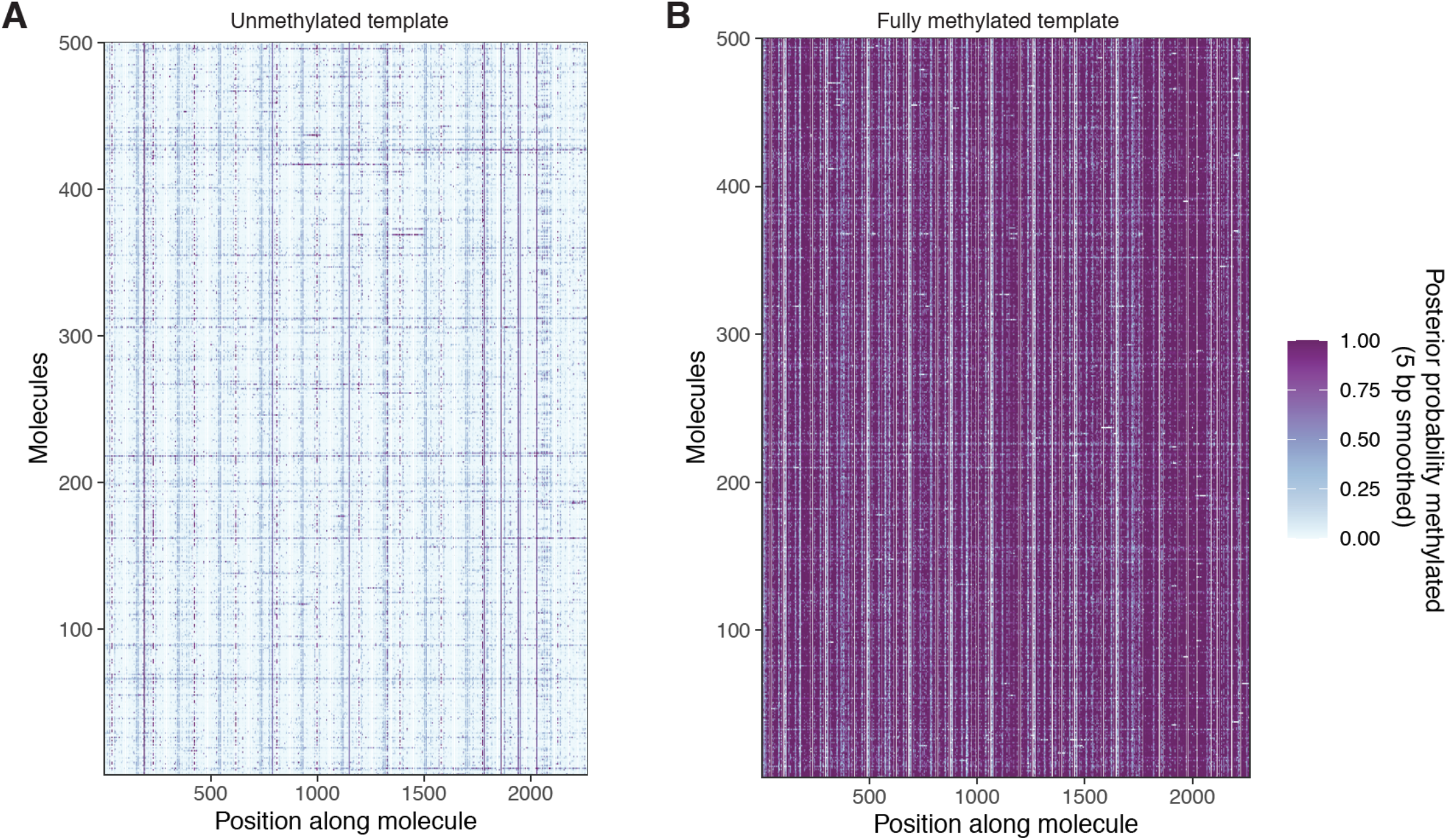
Unmethylated and fully methylated array DNA does not display the same periodic patterning of modified bases seen in methylated chromatin. **A.)** Smoothed modification probabilities for 500 molecules of unmethylated array DNA. **B.)** Smoothed modification probabilities for 500 molecules of fully methylated naked array DNA. In both cases, data are smoothed using a 5 bp rolling mean on the calculated modification probabilities at template A nucleotides.

**Supplementary Figure 6:**
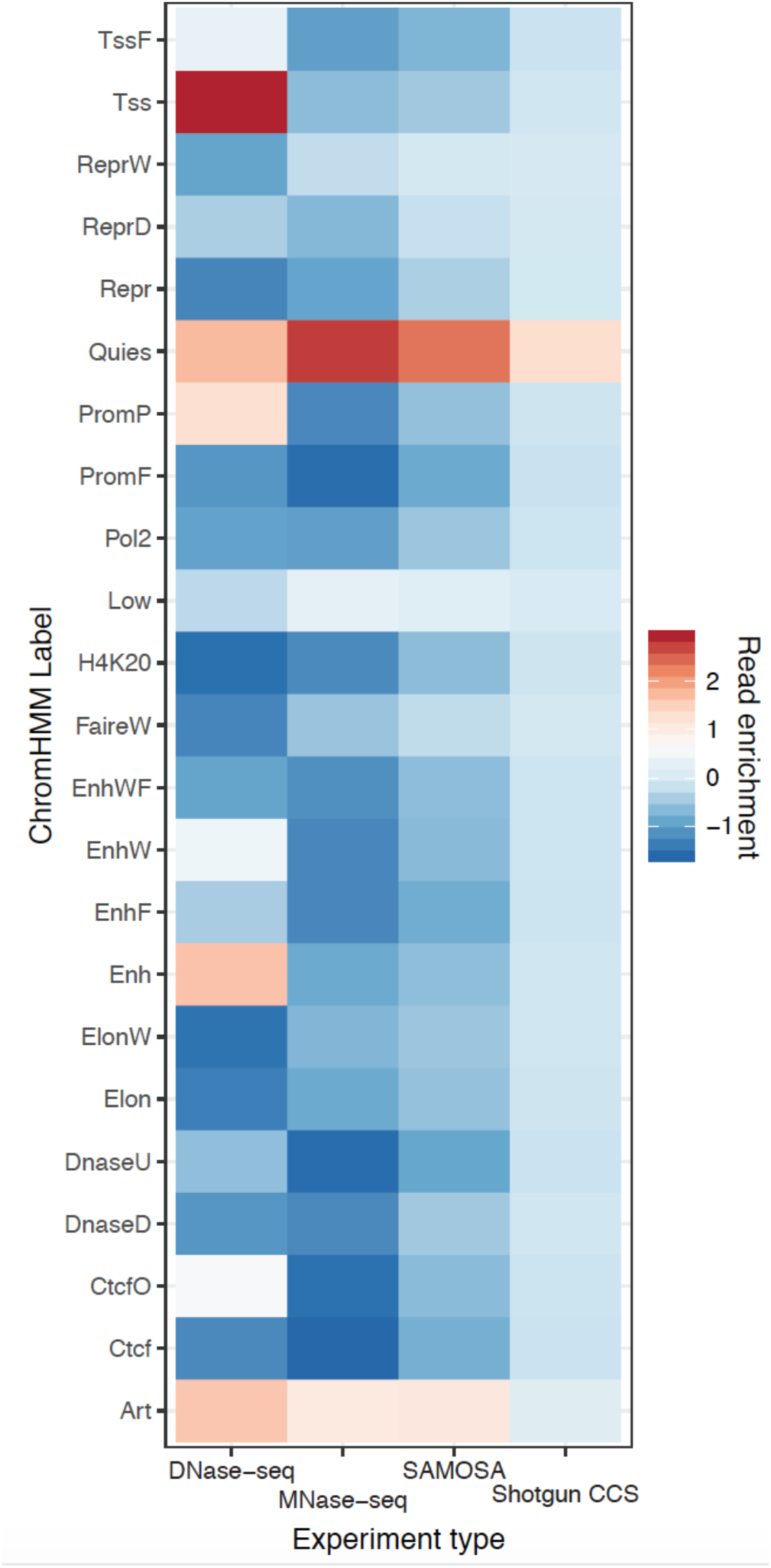
Coverage enrichment of SAMOSA versus other data types at ChromHMM annotated genomic regions. SAMOSA coverage is less biased for open / active chromatin than short-read assays of chromatin accessibility, and is more comparable to short-read MNase-seq assays. SAMOSA coverage is more biased than shotgun PacBio sequencing of the human genome, likely due to the combined use of MNase as a cleavage reagent and the chromatin solubilization protocol used.

**Supplementary Figure 7:**
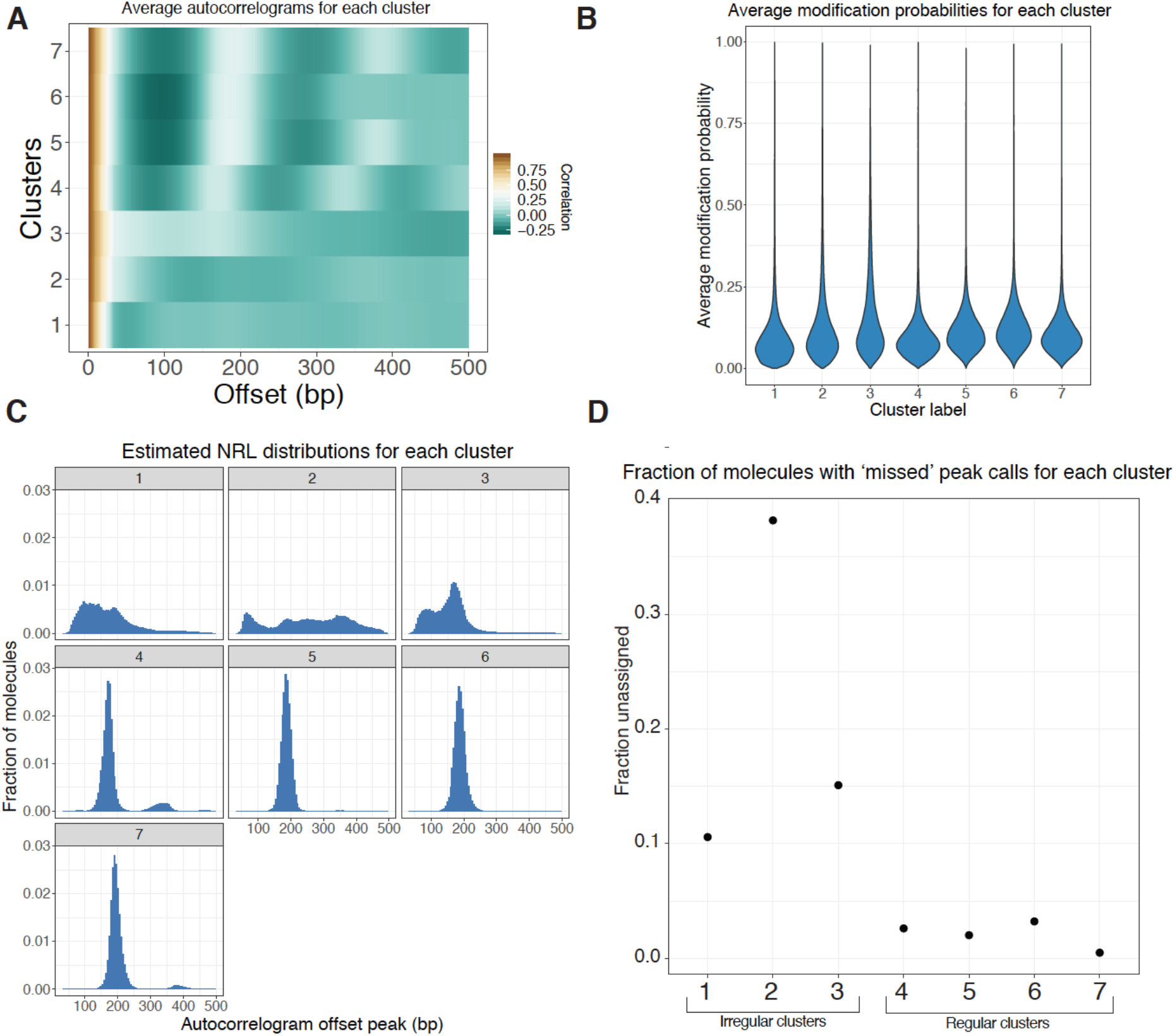
Further characterization of clustered footprinted molecules. **A.)** Average autocorrelograms for the seven Leiden clusters. **B.)** Violin plots of the single-molecule average modification probabilities for each cluster. Clusters do not substantially differ with respect to modification probability, suggesting that clustering is not simply driven by methylation extent. **C.)** As in **Figure 3C**; NRL distribution estimates for each of the seven clusters. **D.)** Autocorrelogram peak-calling fails in a fraction of reads in each cluster; this fraction appears to be negatively associated with the ‘regularity’ of the cluster.

**Supplementary Figure 8:**
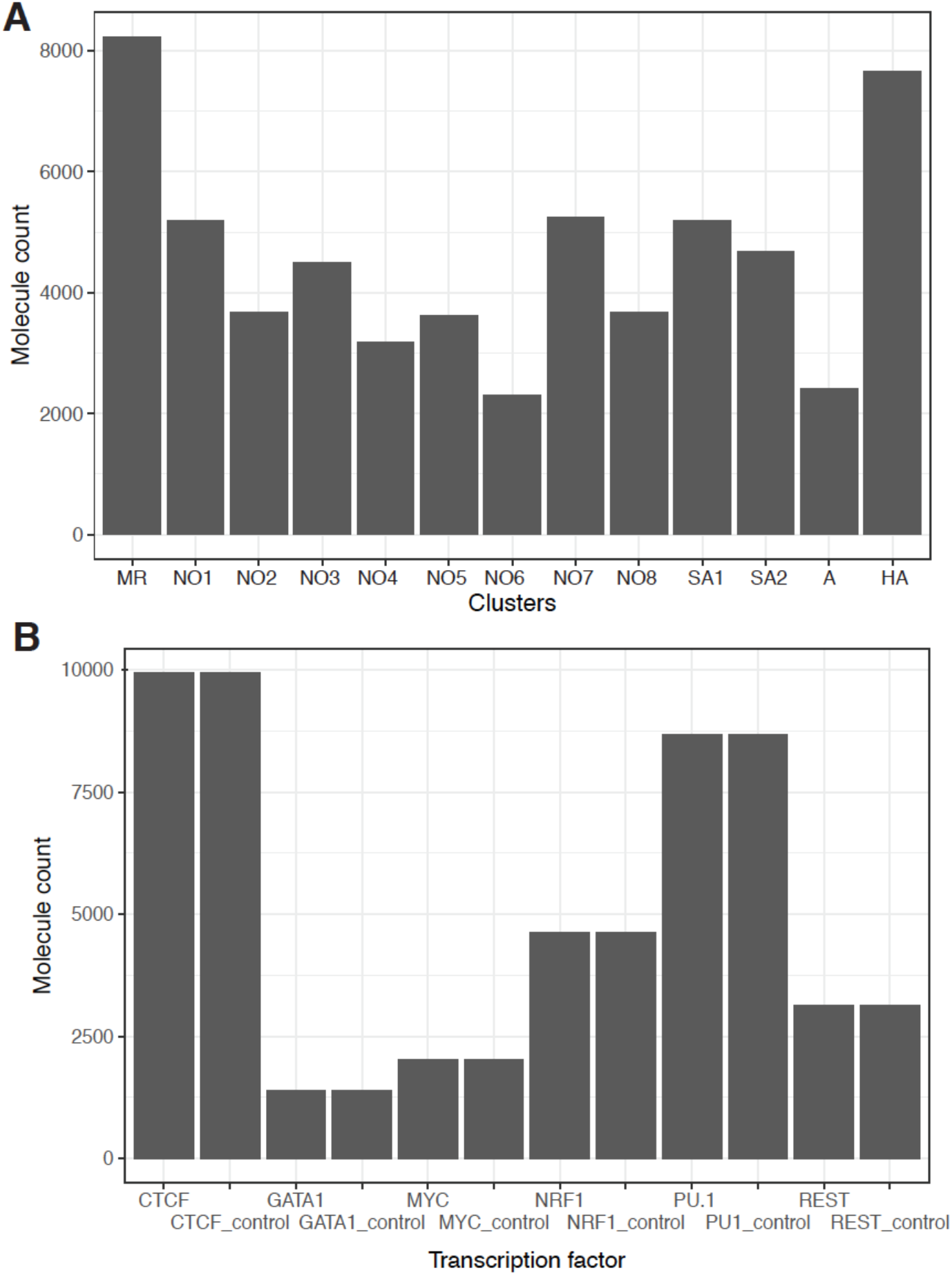
Cluster sizes and numbers of motif-containing molecules for each transcription factor chosen for study. **A.)** Leiden cluster sizes for cluster shown in Figure 4. **B.)** Counts of molecules harboring respective transcription factor binding sites. For each transcription factor, we sampled an equal number of randomly drawn molecules taken from regions GC- / repeat-content matched against TF ChIP-seq peaks.

**Supplementary Figure 9:**
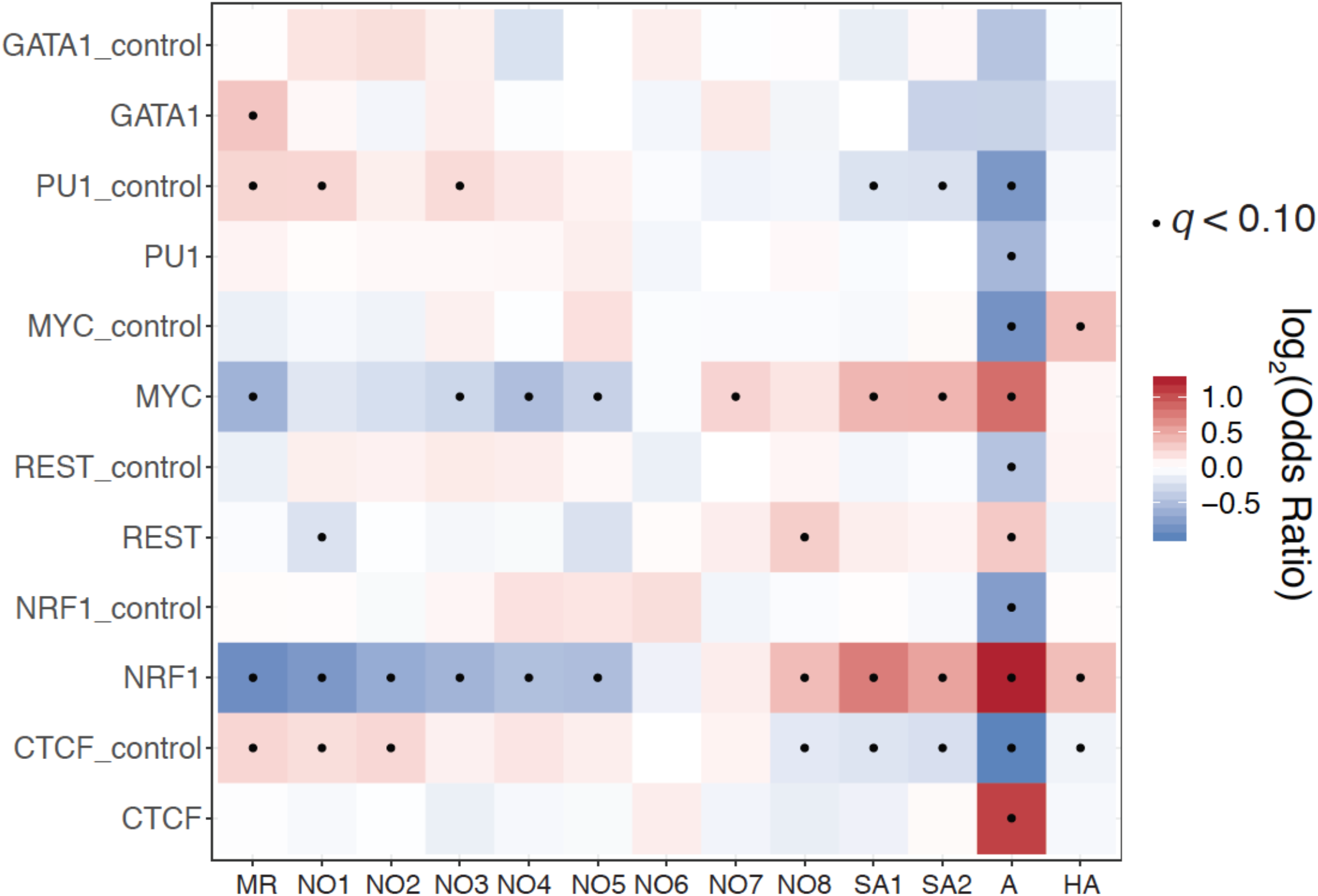
Cluster enrichment for each transcription factor studied. We performed Fisher’s exact tests to determine relative enrichment and depletion of each cluster for each transcription factor surveyed in **Figure 4.** Cluster ‘A’ is consistently depleted across control molecules but enriched across molecules containing bona fide transcription factor binding motifs, suggesting that the clusters identified in this study are biologically relevant. Fishers Exact test odds ratios are plotted in heatmap form and all enrichment tests that are statistically significant under a false discovery rate of 10% (*q* < 0.1) are marked with a black dot.

